# EMoMiS: A Pipeline for Epitope-based Molecular Mimicry Search in Protein Structures with Applications to SARS-CoV-2

**DOI:** 10.1101/2022.02.05.479274

**Authors:** Vitalii Stebliankin, Prabin Baral, Christian Balbin, Janelle Nunez-Castilla, Masrur Sobhan, Trevor Cickovski, Ananda Mohan Mondal, Jessica Siltberg-Liberles, Prem Chapagain, Kalai Mathee, Giri Narasimhan

## Abstract

**Motivation:** Epitope-based molecular mimicry occurs when an antibody cross-reacts with two different antigens due to structural and chemical similarities. Molecular mimicry between proteins from two viruses can lead to beneficial cross-protection when the antibodies produced by exposure to one also react with the other. On the other hand, mimicry between a protein from a pathogen and a human protein can lead to auto-immune disorders if the antibodies resulting from exposure to the virus end up interacting with host proteins. While cross-protection can suggest the possible reuse of vaccines developed for other pathogens, cross-reaction with host proteins may explain side effects. There are no computational tools available to date for a large-scale search of antibody cross-reactivity.

**Results:** We present a comprehensive Epitope-based Molecular Mimicry Search (*EMoMiS*) pipeline for computational molecular mimicry searches. *EMoMiS*, when applied to the SARS-CoV-2 Spike protein, identified eight examples of molecular mimicry with viral and human proteins. These findings provide possible explanations for (a) differential severity of COVID-19 caused by cross-protection due to prior vaccinations and/or exposure to other viruses, and (b) commonly seen COVID-19 side effects such as thrombocytopenia and thrombophilia. Our findings are supported by previously reported research but need validation with laboratory experiments. The developed pipeline is generic and can be applied to find mimicry for novel pathogens. It has applications in improving vaccine design.

**Availability:** The developed Epitope-based Molecular Mimicry Search Pipeline (*EMoMiS*) is available from https://biorg.cs.fiu.edu/emomis/.

## 1 Introduction

Epitope-based molecular mimicry occurs when antibodies cross-react with two different antigens, triggered by the structural similarity and the physicochemical properties at the binding site (Albert and Inman, 1999). Identification of cross-reactivity may explain heterologous immunity when antibodies for a previous infection from an unrelated organism cross-react with newly encountered pathogens (Welsh *et al*., 2010). It can also explain autoimmune disorders such as rheumatoid arthritis when antibodies cross-react with human proteins (Cusick *et al*., 2012).

Limited studies have described a computational molecular mimicry search process. One of the earliest works is a genome-wide BLAST survey to search for parasite-host molecular mimicry (Ludin *et al*., 2011). Many examples of the sequence similarity approach adapted for molecular mimicry searches can be found in the literature. Examples include the study of virulence mechanisms of pathogenic bacteria (Doxey and McConkey, 2013), fungus-plant interactions induced by cross-reactivity (Armijos-Jaramillo *et al*., 2021), and pathogenicity of *Clostridium botulinum* ATCC 3502 to the human host (Bhardwaj *et al*., 2018). Another approach for molecular mimicry search is to look for similar structural motifs (Kristensen *et al*., 2006). A recent tool called *Epitopedia* combined sequence and structural similarity searches for improved scoring of molecular mimicry candidates of known epitopes (Balbin *et al*., 2021). However, sequence and structural similarity of epitopes from two proteins do not guarantee antibody cross-reaction. Non-consecutive amino acids (AA) of the antigenic protein may affect antibody binding, preventing or enhancing its cross-reactivity. To the best of our knowledge, there are no molecular mimicry search tools available that computationally evaluate antibody cross-reactivity.

Deep learning is a promising approach to overcome the major challenges of investigating proteins at a molecular level. The complexity of protein structures and their dynamic nature make the binding energy function highly unstable and difficult to model (Esmaielbeiki *et al*., 2016). Protein docking algorithms may predict the correct “native” binding pose, but the interaction strength is generally poorly predicted (Weng *et al*., 2020). Physics-based simulations can accurately infer the protein interaction trajectory but require substantial computational resources, posing a significant challenge for large-scale mimicry searches. Deep learning can alleviate the high computational cost of protein-protein interactions prediction and improve predictive accuracy (Graves *et al*., 2020; Gainza *et al*., 2020; Wang *et al*., 2020; Pittala and Bailey-Kellogg, 2020). By training a machine learning tool with sufficient positive and negative examples of binding interface regions, the model learns to estimate the strength of interactions between queried antibody-antigen pairs.

Recent data-driven studies have suggested that non-COVID-19 vaccines may provide partial immunity against the SARS-CoV-2 virus (Pawlowski *et al*., 2021; Rivas *et al*., 2021). Mannar *et al*. experimentally confirmed that several SARS-CoV-2-induced antibodies cross-react with proteins from other viruses such as HIV-1 (Mannar *et al*., 2021) and Dengue virus (Nath *et al*., 2020). The discovery of heterologous immunity resulting from mimicry may suggest using other vaccines for partial protection against a rampant pathogen. Mimicry can also explain how the history of prior infections can provide heterologous immunity against a pathogen. Molecular mimicry can also explain several unexpected side effects of the SARS-CoV-2 infection. We recently reported that molecular mimicry between SARS-CoV-2 Spike and human thrombopoietin (TPO) might induce thrombocytopenia in infected subjects (Nunez-Castilla *et al*., 2021).

In this study, we developed a comprehensive computational Epitope-based Molecular Mimicry Search (*EMoMiS*) pipeline that includes sequence and structural similarity searches followed by deep learning for antibody-antigen binding assessments. We applied it to the SARS-CoV-2 Spike protein. We report the discovery of many potential antibody-antigen cross-reactions and discuss their implications.

## 2 Methods

In this study, a comprehensive computational pipeline called *EMoMiS* was developed (Fig. 1). The method identifies known antibodies that can cross-react with target antigens. After searching the database of antigens for regions of sequence and structural similarity with the target protein, a pre-trained deep learning model is used to evaluate if antibodies, known to recognize the database antigens, can cross-react with the target structure. For this discussion, the term “target protein” refers to the query protein (such as the SARS-CoV-2 Spike) for the search.

**Figure 1.**
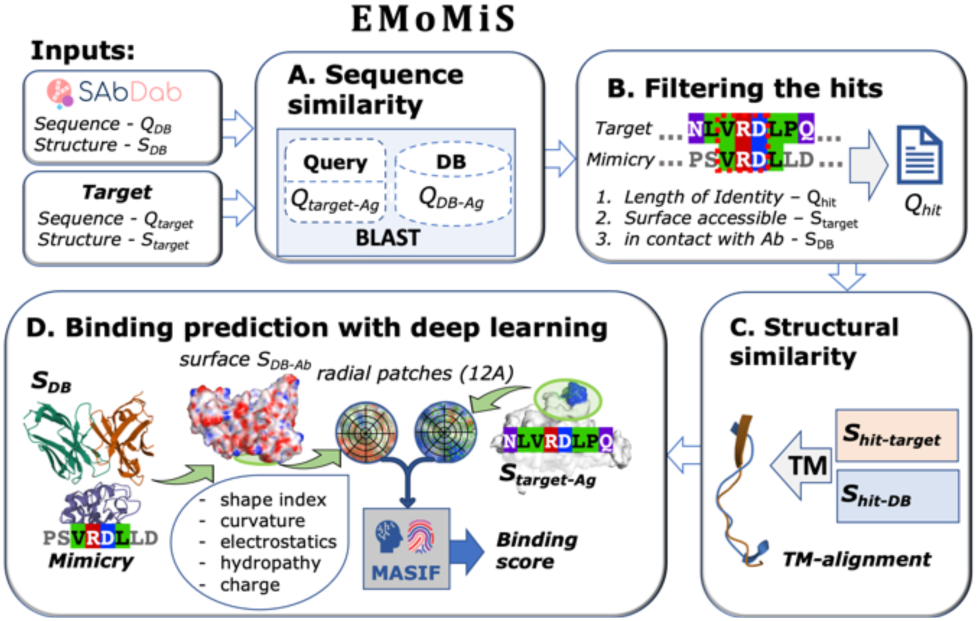
EMoMiS computational pipeline for epitope-based molecular mimicry search. The process consists of four steps: (A) sequence similarity search, (B) filtering, (C) structural alignment, and (D) deep learning for antibody binding evaluation.

The *EMoMiS* pipeline uses the three-dimensional structures, *S*_*DB*_ as well as sequence information, *Q*_*DB*_ of antibody-antigen complexes from the structural antibody database SAbDab (Dunbar *et al*., 2014). The second input to our pipeline is the structure and sequence of the target protein (*S*_*target*_ and *Q*_*target*_), for which mimics are sought. Note that we do not require the availability of a structural complex of the target protein with its antibody. Also, we may have several variant sequences and structures, as is the case with SARS-CoV-2. The target query protein is assumed to be immunogenic, i.e., able to provoke an immune response (Baker *et al*., 2010). In Step A (Fig 1), the target protein sequence(s) (*Q*_*target*_) is searched against the sequences from SAbDab (*Q*_*DB*_) for regions of sequence similarity with other eukaryotic and prokaryotic protein sequences. In Step B, a hit, denoted as *Q*_*hit*_, is retained only if (a) the match has sufficient length (i.e., above a prespecified threshold), (b) the corresponding target antigen is surface-accessible (in *S*_*target*_), and (c) the sequence similarity is in the antibody-antigen interface region (in the corresponding complex in *S*_*DB*_). In Step C, we isolate the target (*S*_*hit*-*target*_) and the mimic (*S*_*hit*-*DB*_) from the hit region and check if they display good local structural alignment. In Step D, we evaluate the potential cross-reactivity using a pre-trained deep learning model (Gainza *et al*., 2020), which estimates the binding strength between the antibody *S*_*DB*-*Ab*,_ complexed with the database antigen *S*_*DB*_ and the target antigen *S*_*target*-*Ag*._.

### 2.1. Sequence Similarity Search

The sequence similarity search between the target protein sequence, *Q*_*target*_, and the antigen sequences from the data set, *Q*_*DB*_, was performed using Protein-Protein BLAST 2.12.0+ (Altschul, 1997). FASTA sequences were downloaded using BioPython utilities (Cock *et al*., 2009). The search was executed with several non-default parameters: the ‘-blastp-short’ flag was used to find small hits because the average epitope length is only about 15 amino acids (Kringelum *et al*., 2013); the ‘-gapopen’ was set to the maximum value since epitope regions are unlikely to allow insertions and deletions without disrupting the binding; and finally, ‘-evalue’ was set to ‘999999’ to include high E-values that often result from short alignments.

### 2.2. Filtering the Sequence Hits

In the filtering step, we select only relevant areas among the hits. First, we require an alignment with an exact match of at least three consecutive amino acids. Second, we check if the three matching amino acids are surface accessible in the target antigen (*S*_*target*_) to allow binding. An amino acid residue is considered surface accessible if its relative accessible surface area (RASA) is more than 20% (Touw *et al*., 2015). Third, we retain hits that lie in the contact region of the antibody-antigen interface in the complex, *S*_*DB*_. An antigen residue is considered a contact point if the distance from any of its heavy atoms (atoms other than H atoms) to the antibody is less than 5Å.

### 2.3. Structural Alignment

The structures of the target and database proteins from each sequence match were assessed for similarity using TM-align, an algorithm for sequence-independent structural comparisons (Zhang, 2005).

To distinguish between molecular mimicry patterns and coincidental structural alignments, we obtained a distribution of alignment scores of randomly selected short motifs. The metric for structural similarity was the root-mean-square deviation (RMSD) between aligned residues in angstroms (Å). Since shorter motifs are expected to have lower RMSD values, the distribution of RMSD values for each motif length was considered separately. For 3000 random antigens from the SAbDab database, we isolated possible epitopes of lengths ranging from 3 to 32 AA, such that the center of each motif was in contact with its native antibody. The isolated motifs of the same length were randomly paired and aligned with the command-line tool, TM-align (Zhang, 2005).

The resulting distribution of structurally aligned random epitopes was used to establish thresholds for acceptable structural mimicry (Fig. S1). The Z-scores were computed for each point of the distribution. RMSD values with Z-score less than -1.645 (one-tailed p-value < 0.05) were considered “high-confidence” hits, while Z-scores between -1.645 and -1.281 (one-tailed p-value < 0.1) were labeled as “medium-confidence” hits (see details in Table S1). Since the epitope is expected to be 32 AA (Kringelum *et al*., 2013), we chopped longer sequence matches into 32 residue-long motifs centered at the point of antibody contact.

### 2.4. Binding Prediction with Deep Learning

After determining the sequence and structure similarity of a candidate binding site along with its surface accessibility, the next step in the *EMoMiS* pipeline is to evaluate the strength of binding between the antibodies of the mimicking proteins to the target protein at the candidate site (Fig. 1). The Molecular Surface Interaction Fingerprint Search (MaSIF-Search) that uses a geometric deep learning approach was used to evaluate antibody-antigen binding, (Gainza *et al*., 2020). The pre-trained model from MaSIF-Search was used for this work. The MaSIF method is designed to uncover patterns on the surfaces of proteins. Given two surface regions (patches) from distinct proteins, the model evaluates compatibility for forming a stable binding complex. The patches are obtained by drawing a fixed-sized geodesic radius on the solvent-excluded protein surface (Sanner *et al*., 1996). The data structure of the patch is a grid of 80 bins with five angular and 16 radial coordinates. Each bin has five geometric and chemical features: shape index, distance-dependent curvature, electrostatics, charge, and hydropathy. The model captures the general distribution of native binding versus decoys by training the artificial intelligence (AI) network with many examples of binding and non-binding protein surface regions.

In Step D of the *EMoMiS* pipeline, we evaluate the structurally similar motifs for the binding strength of the antibody-antigen pair. The antibody patch corresponding to the mimicry antigen is centered on the antibody contact residue. The antigen patch of the target protein (Spike) is centered on the middle amino acid from the filtered sub-query. The resulting patch pair is passed to the pre-trained MaSIF-Search deep learning model that outputs a binding score. If several patches map to the same residue, then we report the binding score of the best patch pair.

The patch extraction method was adapted from the original MaSIF-Search described in (Gainza *et al*., 2020). First, we triangulated each protein complex with a granularity of 1 Å. Geometrical and chemical features for each point in a surface mesh were computed with the MaSIF data preparation module. Next, the structures were discretized into a set of overlapped patches with a radius of 12 Å, where each point in a surface mesh is treated as a patch center.

To evaluate the complementarity of the two patches, the pre-trained MaSIF-Search “sc05” model with a patch radius of 12 Å was used (Gainza *et al*., 2020). The model takes a single patch as input and embeds it into an 80-dimensional descriptor. The original model was trained to minimize the distance between embedded vectors from native binders (positive) and maximize it for non-interacting decoy patches (negative) (Gainza *et al*., 2020). The distance between embedded vectors from two patches will be referred to as the Deep Learning binding score or “DL score.”

### 2.5. Determining Thresholds for Binding Strengths

The pre-trained MaSIF model “sc05” performed well on the antibody-antigen complexes with the receiver operator characteristic area under the curve (ROC AUC) 95% (Fig. 2). Although the “sc05” model was trained on a general collection of protein-protein interactions, we tested the model only on antibody complexes. The structures from 433 non-redundant SAbDab complexes with 90% maximum sequence identity were extracted for the model testing (Dunbar *et al*., 2014). Antigens homologous to any protein from the MaSIF-Search training set were excluded from the testing set, which resulted in 179 protein complexes. Two antigens were called homologous if pairwise alignment identity were greater than 95% (Rice *et al*., 2000). Eighteen Spike-antibody complexes were tested separately, as the SARS-CoV-2 virus was our primary focus. A patch pair from two proteins were labeled “positive” if the distance between patch centers is less than 1Å, while “negative” non-interactive patches were chosen randomly. We observe that the MaSIF pre-trained model is able to distinguish between positive native binders (Fig. 2, orange) and negative decoys (Fig. 2, blue) with a ROC AUC of 95%. The binding scores for SARS-CoV-2 and antibodies (Fig. 2, right) had a similar pattern as the general antibody-antigen complexes from SAbDab (Fig. 2, left).

**Figure 2.**
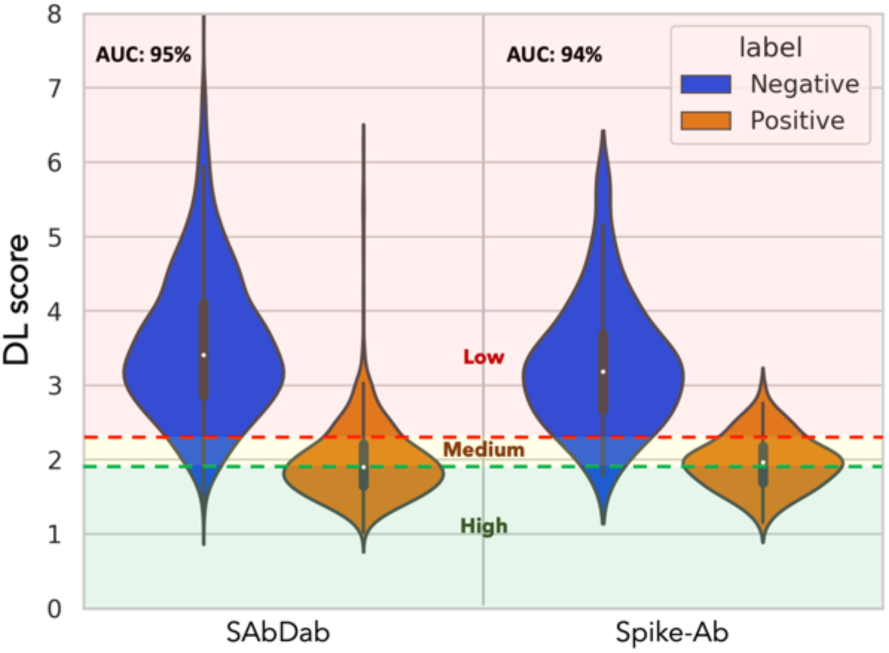
Distribution of Deep Learning (DL) scores of MaSIF pre-trained “sc05” model applied to antibody-antigen complexes from the SAbDab database (left) and the SARS-CoV-2 Spike proteins complexes with native antibodies (right).

Z-statistics for the DL scores distribution for negative non-binding patches (Fig. 2, blue, left) were used to determine the containment thresholds. The value corresponding to a Z-score of -1.645 (one-tailed p-value=0.05) was used as the high-confidence threshold (Fig 2, green). The scores corresponding to one-tailed p-values between 0.05 and 0.1 (Z-scores less than -1.281) were considered a medium-confidence binding (Fig. 2, yellow). The rest of the scores were “low-confidence” (Fig 2, red).

### 2.6. *EMoMiS* Reversed Phase

If the sequence and structure of the antibodies for the target protein are also available, then an additional phase called the ‘reverse’ phase may be added to the pipeline. The pipeline follows the same procedure while the lists of database and target proteins are switched. In other words, every location of the database proteins is queried against the list of known epitopes from the target protein. Such a trick allows searching for cross-reactivity of antibodies, known to recognize the target virus, with the database antigens. When native antibody structures are not available, such as what may happen in the early stages of an epidemic, the pipeline can only be executed in the forward phase. In the case of SARS-CoV-2, such antibodies and their structures are indeed available.

### 2.7. Hardware

The *EMoMiS* software was built in the Chameleon Cloud environment on CHI@TACC “Haswell” bare metal instance, which had 2x 12 core Intel Xeon E5-2670 v3 and 128 GB of RAM (Keahey *et al*.). To ensure reproducibility, the running environment was containerized with Singularity version 3.8.5 (Kurtzer *et al*., 2017). Deep Learning models were tested on a server machine at the Knight Foundation School of Computing and Information Sciences at Florida International University, which had 8 GeForce GTX 1080 Ti GPU, 256G of RAM, and 28 core Intel(R) Xeon(R) CPU E5-2650.

## 3 Results

An epitope-based molecular mimicry search (*EMoMiS*) pipeline was developed in this work. The process consists of four steps: sequence similarity search, filtering, structural alignment and filtering, and deep learning for antibody binding evaluation (Fig. 1). The developed software was applied to search for epitopes from multiple organisms that may mimic a portion of the SARS-CoV-2 Spike protein surface.

### 3.1 Dataset

Structural antibody database SAbDab was used as the reference for the molecular mimicry search pipeline (Dunbar *et al*., 2014). The database was downloaded on November 30, 2021, containing 3,670 unique antibody-antigen complexes. The target sequences and structures of 167 SARS-CoV-2 spike protein references were isolated from the database. The target proteins of homologous MERS and SARS virus families were excluded from the database resulting in 3,193 reference antibody-antigen complexes (Supplementary Table S2).

The Beta, Delta, and Omicron variants of SARS-CoV-2 Spike were obtained from Protein Data Bank (PDB) database (PDB IDs 7LYM, 7V7N, 7T9J) (Gobeil *et al*., 2021; Berman *et al*., 2000).

### 3.2 *EMoMiS* Forward Phase

During the forward phase of the *EMoMiS* pipeline, every location of the 167 target SARS-CoV-2 Spike proteins was matched against antibodies from 3,193 database complexes. The method identified four molecular mimicry candidates, where antibodies from other organisms that likely cross-react with the SARS-CoV-2 Spike (Table 1, A-D). (A) The TN1 antibody from human thrombopoietin (Feese *et al*., 2004) had the most significant Deep Learning score (DL score = 1.281, p-value = 0.01), while structural similarity had medium confidence (RMSD=0.9, p-value=0.093). The mimicry is predicted for the TQLPP motif in the Spike N-terminal domain (NTD). (B) An antibody from tumor necrosis factor receptor superfamily member 5 (TNFRSF-5) (Argiriadi *et al*., 2019) also had high confidence to cross-react with Spike in the NTD region (DL score = 1.749, p-value = 0.032), while the motif ESEF structurally aligned with medium confidence (RMSD = 0.42, p-value = 0.095). (C) The motif NITN from human ABCB-1 (Alam *et al*., 2018) had medium structural similarity with the SARS-CoV-2 Spike protein in the receptor-binding domain (RBD), whereas human-specific inhibitory antibody UIC2 showed a medium score for cross-reaction with the Spike protein (DL score = 2.259, p-value = 0.092). (D) The scores for antibody 2B7 from Dengue virus (Biering *et al*., 2021) to cross-react with Spike were close to medium (DL score = 2.347, p-value = 0.108), while structural similarity for the corresponding NLVK motif had a medium RMSD score (RMSD = 0.25, p-value = 0.061).

**Table 1.**
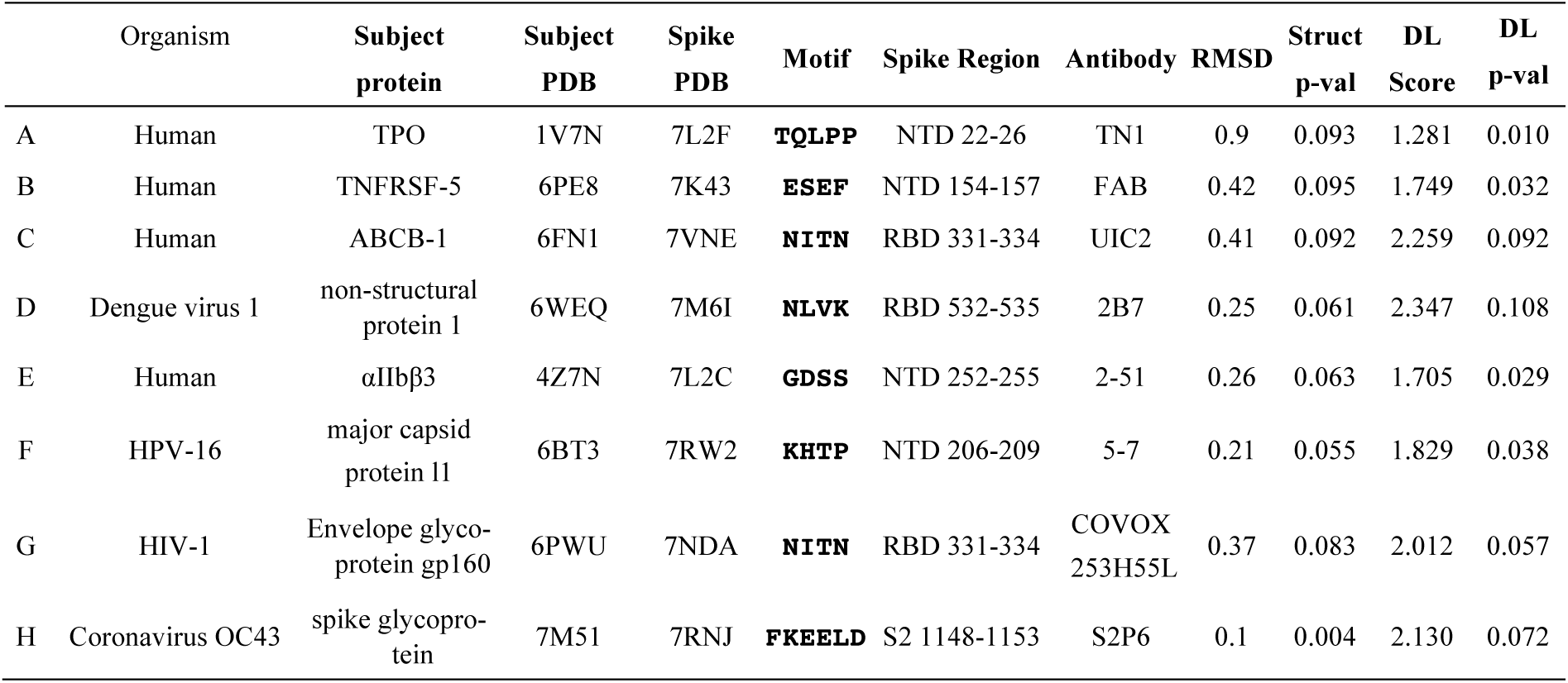
Molecular mimicry for SARS-CoV-2 Spike protein predicted by the *EMoMiS* pipeline. (A-D) Forward phase - antibodies from other organisms predicted to cross-react with Spike. (E-H) Reverse phase - antibodies originated from Spike predicted to cross-react with antigens from the database.

### 3.3 *EMoMiS* Reverse Phase

In the reverse phase of the *EMoMiS* pipeline, the database and target lists were switched to identify unknown epitopes from multiple organisms that may cross-react with native anti-Spike antibodies (Table 1) *EMoMiS* predicted that SARS-CoV-2-specific antibodies can cross-react with (E) human αIIbβ3 (Lin *et al*., 2016), (F) major capsid protein l1 of Human papilloma virus (HPV) 16 (Guan *et al*., 2017), (G) HIV-1 envelope glycoprotein (Pan *et al*., 2020), and (H) coronavirus OC43 Spike glycoprotein (Sauer *et al*., 2021). Human αIIbβ3 motif GDSS and HPV-16 motif KHTP had a high score for antibody cross-reaction (DL score = 1.705 and 1.829, p-value < 0.05), while structural similarity with the same motifs of Spike protein had a medium score (RMSD = 0.26 and 0.21, p-value < 0.1). The NITN motif of the HIV-1 envelope glycoprotein gp160 and SARS-CoV-2 spike protein had medium structural similarity (RMSD = 0.37, p-value = 0.083), while the binding of antibody COVOX-253H55L with the motif of HIV-1 had medium confidence (DL score = 2.012, p-value = 0.057). The FKEELD motif from Spike Coronavirus OC43 had high structural similarity with SARS-CoV-2 Spike protein (RMSD = 0.1, p-value = 0.004), while the binding of S2P6 antibody had only medium confidence for antigen cross-reaction (DL score = 2.130, p-value = 0.072).

### 3.4 Surface Accessibility of Predicted Mimicry Candidates

All predicted mimicry motifs appeared surface accessible in the Spike protein (Fig. 3), confirming the possibility for antibody cross-reaction. Motifs GDSS (Fig. 3, E), KHTP (Fig. 3, F), NITN (Fig. 3, C and G), and FKEELD (Fig. 3, H) are confirmed antibody interacting spots on Spike (Cerutti, Guo, Zhou, *et al*., 2021; Cerutti, Guo, Wang, *et al*., 2021; Dejnirattisai *et al*., 2021; Pinto *et al*., 2021). We note that motif FKEELD (residues 1148-1153) is hidden in Fig. 3 H because the structure for residues after 1147 is not available for the full Spike (Cerutti, Guo, Zhou, *et al*., 2021).

**Figure 3.**
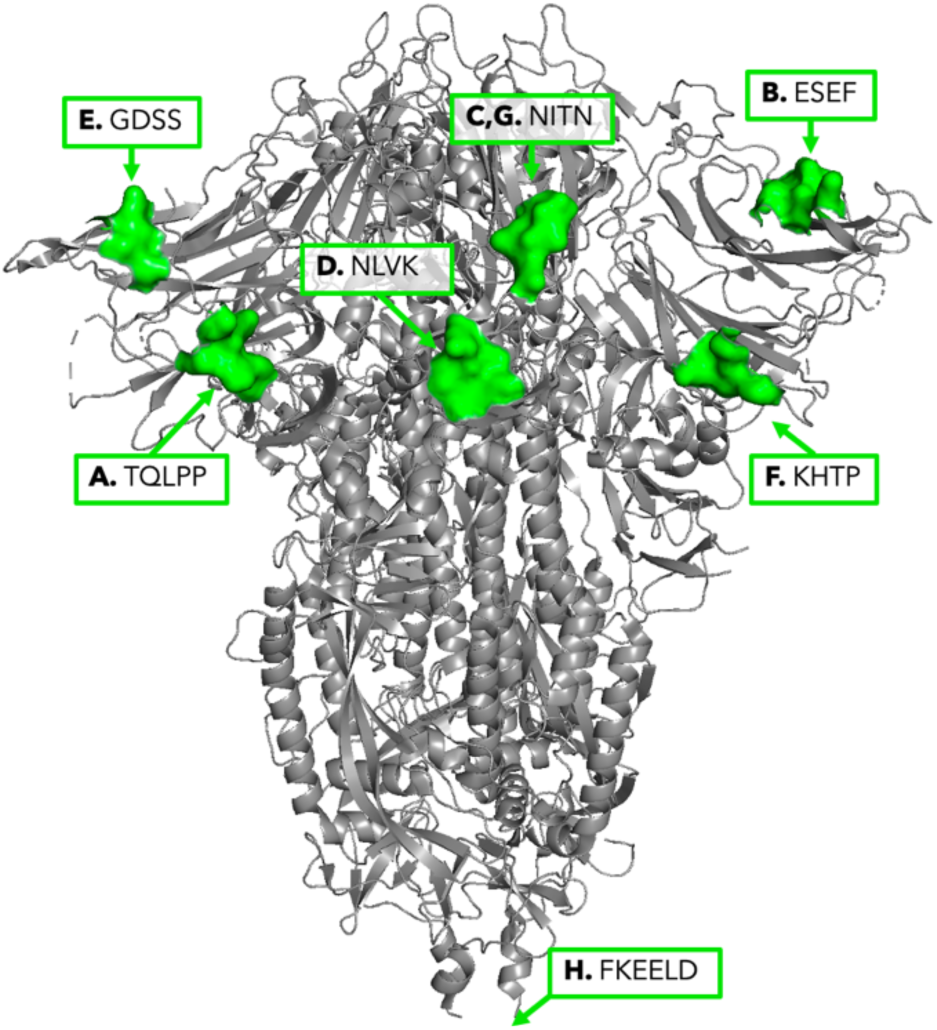
All predicted molecular mimicry motifs on SARS-CoV-2 Spike, identified by the EMoMiS forward and reverse phases.

### 3.5 Impact of Variants

Finally, the SARS-CoV-2 Spike variants of concern (Omicron, Delta, and Beta) were analyzed for the prevalence of identified molecular mimicry candidates (Table 2). The variant Spike structures were queried against the epitopes of four mimicry proteins identified in the *EMoMiS* forward phase: TPO, TNFRSF-5, ABCB-1, and non-structural protein 1 of Dengue virus 1. The binding and structural alignment scores for the Alpha variant corresponds to the previously reported molecular mimicry results (Table 1, rows A-D). The scores for Beta variant 1.351 were not computed for TQLPP and ESEF motifs, as the structure is not available for those regions (PDB ID 7LYM). Two mutations are known in the ESEF motif of the Delta B.1.617.2 variant (E156G and F157V), which resulted in a significant decrease in binding strength of FAB and Delta Spike compared to the Alpha variant (DL score 2.87 vs 1.74). No other mutations directly affected the mimicry motifs, yet, the pipeline shows variability in the binding and structural alignment scores.

**Table 2.**
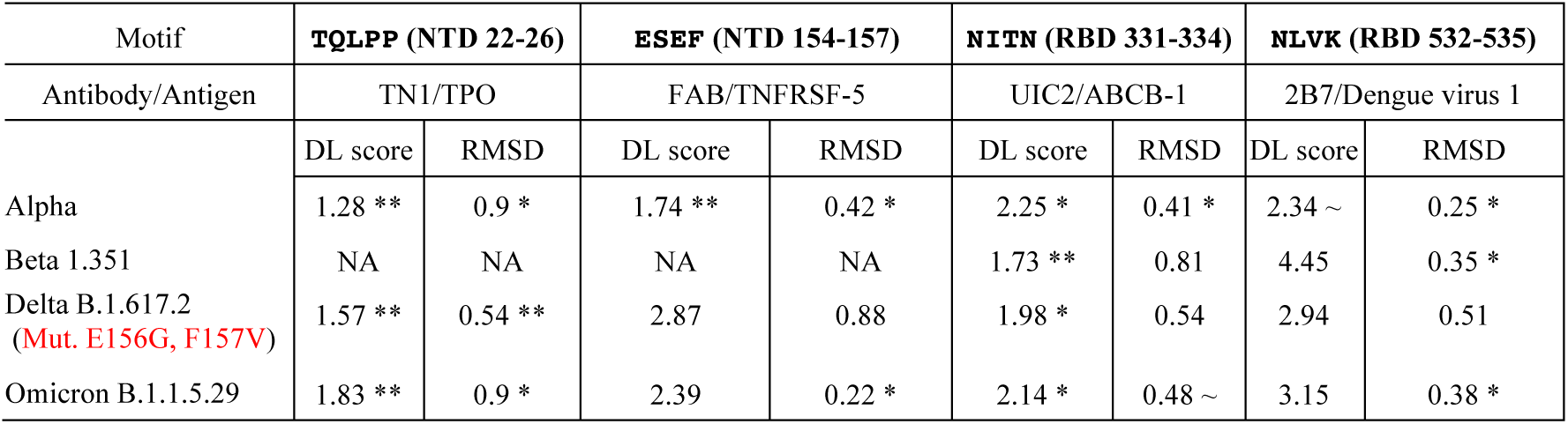
Molecular mimicry predictions across Spike variants. DL score shows the binding strength of the antibody from other organisms (indicated in row 2) to bind the Spike of the given variant. RMSD indicates the structural alignment score of the mimicry antigen and corresponding Spike motifs. DL is denoted as the deep learning score of the antibody (row 2) to bind the N-terminal or receptor-binding domains (NTD and RBD, respectively) of the SARS-CoV-2 Spike. The significance was determined with one-tailed Z-score statistics (∼ p-values<0.11; * p-values<0.1; **p-values<0.05)

The scores for molecular mimicry in the TQLPP region were consistently significant across the variants (Table 2, column 2). The binding score of cross-reacting antibody with ESEF motif significantly decreased in Delta and Omicron variants (Table 2, Column 3, DL score), while its structures in the TNFRSF-5 and Omicron Spike aligned with the same confidence (Table 2, column 3, RMSD). The confidence of the cross-reactive binding strength increased for the NITN motif in Beta 1.351 while staying the same for other Delta and Omicron (Table 2, column 4, DL score). On the other hand, the RMSD between NITN aligned residues increased significantly for every Spike variant (Table 2, column 4, RMSD). The DL score of the 2B7 antibody to bind to Spike has decreased in every variant compared to control (Table 2, column 5, DL score). However, the structural alignment showed a significant decline only in the Delta B.1.6.17.2 (Table 2, column 5, RMSD).

## 4 Discussion

We developed an epitope-based molecular mimicry search pipeline that identifies epitopes that can elicit antibodies cross-reactive to the surface-accessible query viral immunogens. Other molecular mimicry search studies have been fairly simplistic and have not been viewed as an instrument to prepare for the viral pandemic. After applying the *EMoMiS* pipeline to the SARS-CoV-2 Spike protein we hypothesize how cross-reactivity can impact immune response.

### 4.1. *EMoMiS* Pipeline

The *EMoMiS* tool has many advantages over alternative molecular mimicry search methods. First, along with the standard sequence and structural similarity searches, our method evaluates cross-reactive antibody binding strength. The sensitivity analysis revealed that the MaSIF deep learning model accurately separates positive and negative binding with AUC equal to 95% (Fig. 2). Combined with sequence and structural similarity filtering, the deep learning model aims to select only relevant candidates for antibody cross-reaction. Second, our method is capable of predicting cross-reaction with unknown epitopes. Only one protein requires an antibody binding in the sequence similarity region, while the evaluated motif on the second protein can be in any surface accessible position. Thus, the usage of structural antibody database SAbDab expands the search space when compared to the Immune Epitope Database (IEDB), typically used to search for mimicking epitopes (Vita *et al*., 2019). Third, the pipeline may take as input multiple structural conformations and target protein variants, which accounts for protein dynamics. Large variability of target protein structures increases the chance of finding configuration favorable for antibody binding.

Yet, despite the high accuracy, the deep learning model for cross-reactivity evaluation needs improvements. The model can be biased towards some interface structures. The training and testing sets for MaSIF deep learning are limited by the protein complexes available in the Protein Data Bank (PDB). If the target protein is sufficiently novel in comparison to the contents of the database and the training set, the model may fail to generalize and may produce false predictions. Another limitation of the deep learning model is the absence of glycans in the set of features. The glycosylation events may significantly affect the antibody neutralizing properties and thus, the model sensitivity (Miranda *et al*., 2007). Those limiting factors may consolidate the false-negative predictions. For example, the score for the antibody from the Ab-Spike OC43 complex (PDB ID 7M51) to bind SARS-CoV-2 Spike protein (PDB ID 7M53) was very low (DL score = 3.066, p-value = 0.306, see Supplemental Table S3). Yet, the cross-reaction between antibody B6 (PDB IDs 7M51 and 7M53) with coronavirus OC43 and SARS-CoV-2 Spike proteins was experimentally verified (Sauer *et al*., 2021). On the other hand, another Spike configuration (PDB ID 7RNJ) was found to cross-react with coronavirus OC43 (Table 1, H), which highlights the advantage of using multiple target structures.

While the lack of a molecular structure for the target protein may be seen as a limitation for the *EMoMiS* pipeline, protein structures can be quickly and accurately predicted with the recent advances in the deep learning field (Jumper *et al*., 2021).

### 4.2. Antiviral Antibody Cross-reaction

The *EMoMiS* pipeline identified four sites in the SARS-CoV-2 Spike protein that may mimic epitopes from other viruses. Antibody cross-reaction with viral mimic epitopes may provide cross-protection to the host. On the other hand, antibodies elicited by mimicry may cross-react with the vital antigen with lower affinity. As a result, bound antibodies may fail to block the virus from cell entry, while shielding the pathogen from its native antibodies. Such an effect is known as antibody-dependent enhancement (ADE), where antibody enhances the viral entry (Tirado and Yoon, 2003).

The strongest antibody cross-reactions were predicted for HPV-16 major capsid protein l and the SARS-CoV-2 Spike protein. HPV is a DNA virus that can cause benign and malignant neoplasms (Molijn *et al*., 2005). A recent study reported a patient for which persistent verruca vulgaris (benign HPV warts) paradoxically regressed after recovery from COVID-19 (Demirbaş *et al*., 2021). However, we hypothesize that the HPV vaccine will not be effective against COVID-19, as the predicted motif KHTP is not included in the list of cross-neutralizing epitopes (Tumban *et al*., 2011). Indeed, a recent study of immunization records shows that there is no decrease in the frequency of COVID-19 cases in HPV vaccinated patients (Pawlowski *et al*., 2021).

The next medium confidence molecular mimicry was predicted for non-structural protein 1 of dengue virus and the SARS-CoV-2 Spike protein (Table 1, row D). The antibody cross-reaction between proteins from the dengue and SARS-CoV-2 viruses was previously computationally predicted (Nath *et al*., 2020). A recent study experimentally confirmed this hypothesis, but the rate of antibody cross-reaction was only 22% (Lustig *et al*., 2021). Yet, it remains unknown if antibodies induced by such cross-reaction can provide cross-protection.

Another molecular mimicry hit was found between the SARS-CoV-2 Spike and the HIV-1 envelope glycoprotein. The first hint to cross-reactivity was documented in the study that observed false-positive HIV tests in COVID-19 patients (Tan *et al*., 2021). Another group has found that two out of nine tested anti-HIV antibodies can cross-react with the Spike protein, but such antibody reactions do not block viral entry (Mannar *et al*., 2021). Thus, cross-reaction of anti-HIV antibodies with Spike may promote an adverse effect of antibody-dependent enhancement (ADE).

The last molecular mimicry prediction was for coronavirus OC43 and SARS-CoV-2 Spike proteins. The structure of the predicted cross-reactive anti-Spike antibody S2P6 was isolated from the experimental study confirming the existence of such mimicry (Pinto *et al*., 2021). The FKEELD region from perfusion-stabilized S ectodomain trimers is a confirmed epitope and conserved across SARS-CoV, MERS-CoV, SARS-CoV-2, and OC43 (Pinto *et al*., 2021). Identified mimicry proves the validity of our pipeline. We note that SARS-CoV and MERS-CoV are absent in the mimicry results because they were excluded from the database to avoid obvious molecular mimicry from the closely related viral families.

### 4.3. Autoimmune Disorders

Molecular mimicry between epitopes from viral and human proteins may cause autoimmune disorders. Antibodies induced by the virus may bind to essential human proteins and thereby change their ability to function. The SARS-CoV-2 infection can initiate a cascade of interactions between plasmin, complement, and platelet-activating systems, which can lead to tissue damage, thrombosis, inflammation, and cytokine storm (Mukund *et al*., 2020). The nature of the adverse interactions can be caused by the auto-antibodies induced by molecular mimicry. Four of the identified mimicry antigenic motifs originated from human proteins: TPO, TNFRSF-5, ABCB-1, and αIIbβ3 (Table 1).

The strongest cross-reaction was predicted for antibody TN1 to recognize the TQLPP motif from TPO and the Spike protein (row A in Table 1). Human TPO is a glycoprotein hormone that regulates the production of platelets, essential elements for blood coagulation (Kuter and Begley, 2002). Previously, we have shown that molecular mimicry between Spike and TPO may induce thrombocytopenia, a disorder with low blood platelet count (Nunez-Castilla *et al*., 2021). The *EMoMiS* pipeline result provides further evidence that antibodies that recognize TQLPP in Spike may cross-react with TPO, causing the reduction of platelet counts. Thrombocytopenia is a common side effect in COVID-19 patients and is associated with an almost 5-fold increase in mortality (Yang *et al*., 2020; Shi *et al*., 2021).

Another high-confidence molecular mimicry was predicted for SARS-CoV-2 Spike and the human protein TNFRSF-5 (Table 1, row B). TNFSF-5, also known as CD40, is a tumor necrosis factor receptor superfamily member expressed by immune and non-immune cells and involved in producing pro-inflammatory cytokines (Vonderheide and Glennie, 2013). Previous studies have reported that monoclonal antibodies against the CD40 ligand may induce thrombophilia (Kawai *et al*., 2000). At the same time, it has been shown that COVID-19 infection increases susceptibility to systemic thromboembolic complications (Mui *et al*., 2021; Ferrari *et al*., 2020; Oudkerk *et al*., 2020). We hypothesize that thrombophilia in COVID-19 patients can be induced by the antibody cross-reactivity between CD40 and Spike.

The next high-confidence cross-reactivity prediction was between an antibody from SARS-CoV-2 Spike and human αIIbβ3 (Table 1, E). The protein αIIbβ3 is a heterodimeric platelet receptor that plays an essential role in platelet aggregation (Ma *et al*., 2007). A previous study showed that the level of αIIbβ3 activation on platelets from non-surviving COVID-19 patients decreased compared to survivors (Ercan *et al*., 2021, 5). The mechanism for the decline in αIIbβ3 in severe COVID-19 patients was unknown. Here, we propose that the imbalance of αIIbβ3 can be explained by auto-antibodies induced by Spike.

The cross-reactivity of antibody UIC2 with human ABCB-1 and SARS-COV-2 Spike protein was predicted with medium confidence (Table 1, row C). The multidrug transporter ABCB1 is an ATP-binding cassette transporter that is involved in protecting tissues from toxic insult and plays a role in multidrug extrusion from cancer cells (Alam *et al*., 2018). The implications of this mimicry discovery remain to be understood and would require data from COVID-19 patients with cancer.

### 4.4. Effect of Mutations on Molecular Mimicry

When the structure and sequence of viral variants are available, molecular mimicry results may explain the change in the immune response upon evolutionary mutations. For example, the antibody known to recognize TNFRSF-5 has a reduced cross-reactive binding strength in the Omicron and Delta variants compared to the Spike protein in the reference Alpha strain (Table 2, column 3). These variants have less chance to elicit auto-antibodies against TNFRSF-5, implying a lower chance of developing thrombophilia. On the other hand, TQLPP molecular mimicry of Spike with human TPO was consistent across all SARS-CoV-2 variants (Table 2, column 2), suggesting that thrombocytopenia is a concern of COVID-19 infection regardless of mutation. Additionally, the reduction in binding score between antibody 2B7 and Beta, Delta, and Omicron variants compared to Alpha suggests the potential loss of cross-protection provided by the previous infection of the dengue virus 1.

## 5 Conclusion

In conclusion, we have developed a novel approach to infer epitope-based molecular mimicry. We demonstrate the vital importance of predicting cross-reactivity by applying *EMoMiS* to the SARS-CoV-2 Spike protein. We have found one confirmed mimicry epitope, FKEELD from SARS-CoV-2 and OC43 Spike proteins. Other predicted events of antibody cross-reactivity of SARS-CoV-2 Spike with HPV-16, HIV-1, and Dengue virus were suggested by previous literature (Demirbaş *et al*., 2021; Mannar *et al*., 2021; Lustig *et al*., 2021). Unlike previous studies (Balbin *et al*., 2021; Ludin *et al*., 2011; Doxey and McConkey, 2013), the *EMoMiS* pipeline can predict the exact site of molecular mimicry, thus opening the door for further experimentation. Most importantly, the results should be seen as an attempt to explain observed phenomena in terms of partial immunity and the COVID-19-associated complications and side effects. *EMoMiS* has generated potential explanations for thrombocytopenia and thrombophilia, observed to occur in some COVID-19 patients. All predicted molecular mimicry candidates were derived computationally and should be verified in the laboratory. Additionally, this work serves as a step toward building generic pipelines to prepare for future epidemics caused by new pathogens. Our methods also provide a way to quickly understand what one could expect with new variants of a virus.

## Acknowledgements

The authors thank the members of the Bioinformatics Research Group (BioRG) at FIU for valuable feedback and comments.

