## Supplementary materials for "EMoMiS: A Pipeline for Epitope-based Molecular Mimicry Search in Protein Structures with Applications to SARS-CoV-2"

*This document serves as the supplementary materials for the paper "EMoMiS: A Pipeline for Epitope-based Molecular Mimicry Search in Protein Structures with Applications to SARS-CoV-2". The contents in this document contain supplementary figures and tables to support the claims and conclusions in the paper.*

**Authors**: Vitali Stebliankin[1], Prabin Baral[2] , Christian Balbin[3], Janelle Nunez-Castilla[3], Masrur Sobhan[1] , Trevor Cickovski[1], Ananda Mohan Mondal[1,5], Jessica Siltberg-Liberles[3,5], Prem Chapagain[2,5], Kalai Mathee[4,5], *and* Giri Narasimhan[1,5] *

**Author affiliations:**

[1]Bioinformatics Research Group (BioRG), Knight Foundation School of Computing and Information Sciences, Florida International University, Miami, USA

[2]Department of Physics, College of Arts, Science and Education, Florida International University, Miami, USA

[3]Department of Biological Sciences, College of Arts, Science and Education, Florida International University, Miami, USA

[4]Department of Human and Molecular Genetics, Herbert Wertheim College of Medicine, Florida International University, Miami, USA

[5]Biomolecular Sciences Institute, Florida International University, Miami, USA

### Supplementary Figures

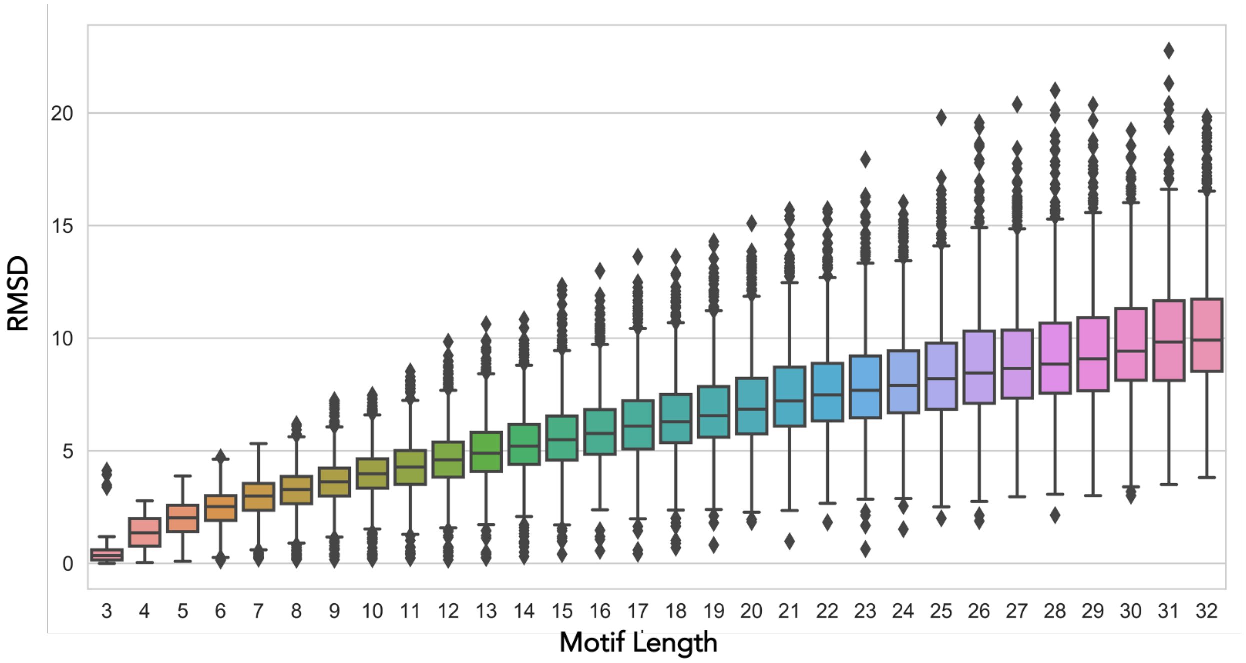


**Figure S1.** Distribution of RMSD values of two randomly aligned motifs of various lengths.

### Supplementary Tables

**Supplementary Table S1**: The thresholds of RMSD values used by EMoMiS to identify the significant structural similarities. The thresholds correspond to Z-score critical values computed from the RMSD random distribution (Fig. S1). The high confidence threshold corresponds to a Z-score of -1.645 (one-tailed p-value<0.05), while RMSD with Z-scores less than -1.281 were considered medium confidence (one-tailed p-value<0.1).

| Motif Length | RMSD high confidence | RMSD medium confidence |
|---|---|---|
| 3 | 0.01 | 0.02 |
| 4 | 0.18 | 0.44 |
| 5 | 0.64 | 0.93 |
| 6 | 1.02 | 1.31 |
| 7 | 1.46 | 1.79 |
| 8 | 1.62 | 1.97 |
| 9 | 1.87 | 2.25 |
| 10 | 2.16 | 2.56 |
| 11 | 2.4 | 2.8 |
| 12 | 2.48 | 2.96 |
| 13 | 2.77 | 3.23 |
| 14 | 2.92 | 3.45 |
| 15 | 3.04 | 3.6 |
| 16 | 3.3 | 3.89 |
| 17 | 3.45 | 4.09 |
| 18 | 3.65 | 4.23 |
| 19 | 3.82 | 4.46 |
| 20 | 3.89 | 4.62 |
| 21 | 4.16 | 4.89 |
| 22 | 4.49 | 5.15 |
| 23 | 4.42 | 5.19 |
| 24 | 4.73 | 5.48 |
| 25 | 4.69 | 5.52 |
| 26 | 4.77 | 5.61 |
| 27 | 4.98 | 5.87 |

| | | |
|---|---|---|
| 28 | 5.11 | 5.99 |
| 29 | 5.38 | 6.3 |
| 30 | 5.75 | 6.6 |
| 31 | 5.83 | 6.73 |
| 32 | 6.08 | 7 |

**Supplementary Table S2.** PDB IDs used by EMoMiS pipeline **A)** SARS-CoV-2 query structures **B)** Database antibody-antigen complexes.

| **A) SARS-CoV-2 Spike PDB IDs** |
| --- |
| 7L06, 7L02, 7DD8, 7DD2, 7K8S, 7CWT, 7DZX, 7ND3, 7D0C, 7M53, 7ND6, 7EJ5, 7LQW, 7K4N, 7ND4, 7D03, 6XCM, 7N9C, 7KXJ, 7L57, 7E3L, 7LSS, 7KML, 7N9T, 7DK6, 7MM0, 7CWM, 7OAN, 7K8W, 7ND8, 6WPS, 7NDB, 7CZT, 7M6G, 7M6H, 7P7A, 7K8T, 7NDC, 7JWB, 6ZHD, 7K8U, 7SO9, 7D0B, 7CZQ, 7CZP, 7R8N, 7L2D, 7KXK, 7KQE, 7NDD, 7KKK, 7B18, 7ND5, 7DK4, 7JW0, 7DZY, 7MY2, 7CZV, 7JV4, 7DCC, 7MJJ, 7LD1, 7N9E, 7DK5, 7LCN, 7D0D, 7N8H, 7K43, 7NTC, 7S0C, 7K8Y, 7P77, 7KSG, 7CZX, 7LAA, 7M6E, 7MJK, 7E5R, 7SOB, 7N0H, 7N0G, 6XEY, 7KMK, 7N5H, 6XCN, 7CWU, 7K8Z, 7CZU, 7JVC, 7M6F, 7VND, 7CAI, 7CWS, 7C2L, 7RW2, 7K8X, 7KQB, 7LXY, 7CZW, 7E3K, 7K8V, 7A29, 7MKL, 7CAC, 7CZS, 7MY3, 7V2A, 7N9B, 7R8M, 6ZXN, 7P79, 7CAK, 7EH5, 7LXZ, 7CZZ, 7LRT, 7VNE, 7K9J, 7P78, 7CHH, 7P40, 7S0D, 7L09, 6WPT, 7DCX, 7E5S, 7E8C, 7L3N, 7KS9, 7KKL, 7LAB, 7V26, 7L2F, 7L58, 7LS9, 6Z43, 7CYP, 7MJH, 7RA8, 7DK7, 7AKD, 7NDA, 7L56, 7R8O, 7LQV, 7LY2, 7D00, 7L2E, 7ND9, 7A25, 7VNC, 7JV6, 7FG2, 7CWN, 7K9H, 7K90, 7EJ4, 7BYR, 7RKV, 7ND7, 7CZR, 7CWL, 6ZDH, 7FG3, 7CZY, 7LJR, 7M6I |
| **B) Database PDB IDs** |
| 7EOW, 7C94, 5DUM, 6PXR, 6WO4, 3UBX, 4NC0, 6MW9, 5XCU, 4HEM, 6PE8, 5WNA, 5TPN, 2DQG, 3IFN, 5DFW, 6NNJ, 7E9B, 7LMP, 6I3Z, 6Z20, 7K39, 2VWE, 6GDG, 6W7S, 4LAJ, 6XBK, 7M30, 7BPK, 1MHP, 6DJP, 3NH7, 6NIJ, 5XCT, 4XMN, 2AEP, 7CMV, 5U3O, 5ACM, 2A0L, 4DAG, 2ZNW, 4O58, 6N5D, 4ZTO, 5F1O, 1EO8, 6HGA, 1KIQ, 7EZM, 6O24, 7KEX, 5CD5, 5VLP, 4YDV, 6A0Z, 7LEZ, 6XPY, 3T3M, 1U95, 5F96, 3UYP, 7K75, 4KHT, 4GMS, 2ITC, 6SC8, 6DKJ, 5F7Y, 3NGB, 2DQC, 3V6Z, 4S1S, 1N6Q, 5U68, 5VXK, 7E33, 3GI9, 7K84, 2NY4, 6K5F, 4HKZ, 5UK0, 3H3P, 4ZXB, 6UC6, 6NI3, 6H72, 6P3S, 5IES, 4ALA, 4NP4, 7RD5, 4XMK, 4WHT, 4KSD, 4XC1, 4M8Q, 3KJ4, 7BC7, 6FYU, 4C2I, 6VRL, 4UU9, 7LMO, 6WJL, 6X8P, 7CMK, 6VJA, 5C3L, 6X58, 6O3B, 4LGS, 3SO3, 5KZP, 6NMU, 6BAE, 4ENE, 5M14, 5U3D, 6WS6, 4D9R, 5LSP, 5BJZ, 5LHN, 4Y8D, 5IF0, 2V17, 5W06, 6BAH, 6C6Y, 6DF2, 2Y36, 4OCM, 6VI1, 7EVM, 5XCQ, 3MLZ, 6OZ6, 6URH, 6J5G, 3HMX, 6APB, 5F21, 5JHL, 6X3Z, 4M7L, 6H6Z, 6ORQ, 6QV0, 5O03, 2Y06, 4R0L, 6R2S, 5W4L, 3U4E, 4UUJ, 6EY6, 6XLZ, 6U38, 6GV1, 2VDR, 6B3J, 4NCO, 4ZS7, 6D0U, 7CEB, 2WUB, 3JBE, 1RVF, 2P4A, 4UIH, 5FGB, 4HHA, 1IC4, 6V80, 6TEJ, 7O31, 6N51, 3VG9, 6OKM, 4XGZ, 1BGX, 6NI2, 2P46, 5V2A, 4O02, 7DKJ, 6N4Y, 2VDN, 1S5H, 2J4W, 3BT2, 4XBE, 6WEX, 7O3B, 3HI1, 6OZ2, 5A2I, 2P49, 1KXV, 3LEX, 1UZ8, 6LXI, 6MQC, |

7NQK, 6JEP, 7CQC, 5A1Z, 6KVF, 6V8X, 6X06, 6Y97, 2VDP, 6X40, 6VC9, 7M2I, 5DMI, 1NMA, 4M1C, 6CXC, 1AR1, 4HLZ, 5U4L, 6MTN, 6Z3P, 6EAY, 7R87, 3MLY, 7ELX, 6LFO, 4WGV, 4Z5R, 5VN8, 6H70, 4W2Q, 3GBM, 6BGT, 7MWX, 6O3C, 6UYD, 5MY4, 4F37, 5N0A, 4RAU, 6FY1, 6MG5, 4WEM, 3EO1, 5IG7, 6B3S, 7C88, 2H2S, 7D3K, 6MLM, 6IEB, 7PIU, 5NHR, 6VUG, 6N50, 6WZK, 6UOE, 5UEM, 5HHX, 5I6X, 6WW2, 7JVQ, 5BV7, 4CAD, 6J7W, 6ULF, 6IAP, 7LF0, 7APJ, 5TBD, 6MHR, 3GB7, 1RJL, 2HMI, 3A6B, 6WEZ, 6K67, 4QY8, 6V7Y, 5ANM, 7NWL, 6X18, 1ZWI, 6QUZ, 5XBM, 6W4M, 6M0F, 6CNK, 6E1K, 7NEZ, 7K78, 2NY3, 6VLR, 6DO1, 5KZC, 5MU2, 1ADQ, 5HI3, 5TSJ, 6WN1, 5EOC, 6LML, 5H32, 6CWG, 6V4P, 3MLU, 3J42, 3G04, 3PGF, 7CEC, 5N7W, 6J71, 4AEI, 5OMM, 6XBM, 3F7V, 6MU7, 5ESV, 6M38, 5W6D, 7KI0, 4QHU, 6XM2, 5E5M, 7M7E, 6QV1, 4DGI, 7D6Y, 3U9U, 6K7O, 7JIX, 4U6G, 6CWT, 6J14, 7D5Q, 5WHK, 3UJI, 4EIZ, 5TH9, 6ADA, 2XXM, 3IFO, 5CIN, 6WN4, 5YOY, 5DA0, 6VTW, 5F3H, 5DWU, 6W0A, 2VYR, 1Z3G, 6B0A, 3WSQ, 4KV5, 6VO3, 6SSP, 7KQ7, 1U8L, 7N4N, 6RQM, 7RSO, 5IVN, 3MLR, 5BO1, 6OZC, 6RUV, 6NIY, 5G64, 6MWC, 4JO3, 3L5W, 7BC6, 4LGP, 6OIJ, 5FXC, 4G6A, 6M3Z, 5T3S, 6VL8, 4PY8, 7LO6, 4N1C, 6VPX, 6PE9, 6JJP, 6DZT, 7NFD, 7RTB, 2X7L, 6K65, 4W6X, 4YXH, 5UM8, 6X7W, 3LHP, 7D3S, 6MAR, 4DGV, 4XWO, 7M7H, 6HUK, 6NC2, 3PJS, 1TZH, 6S5A, 6PZZ, 6W0D, 4N9O, 6I53, 7O06, 4BEL, 5CZX, 6DFJ, 5E8D, 3MUG, 3EFF, 5GGR, 1ZVY, 6UJC, 5MJE, 3LEY, 6WWZ, 4U6H, 5F93, 6IEK, 7EW4, 2VXS, 6X1T, 4LDL, 7DCE, 6EJM, 7CGW, 4GLR, 7RMI, 4XNZ, 5O6V, 2VXT, 6XMI, 6II8, 5XCS, 1V7M, 5YD5, 6LZ4, 5XKU, 4N9G, 4OGY, 5LWF, 6WHA, 4XP4, 2QR0, 7LF7, 4HT1, 2CK0, 5WK3, 6WIZ, 2NXZ, 4LSP, 3S35, 4HCR, 6WXL, 6WDS, 6O9I, 6FGB, 6XR0, 7P14, 5F8Q, 3IFL, 5F97, 5XWD, 3OR7, 6HER, 5WDU, 6WZM, 3BE1, 4I3R, 7LCV, 6XRT, 7JGJ, 1YJD, 1QFW, 3GRW, 7KBI, 6ZTR, 5YWY, 6DCQ, 1U93, 7KEO, 2NY7, 2XZQ, 7P16, 6HJQ, 3LD8, 4R9Y, 5F1K, 3U7Y, 6Z7W, 6W4Y, 6D6U, 1N8Z, 1HEZ, 6S0Y, 3J1S, 6WIO, 5K9K, 7KBT, 7CZ5, 6MXT, 6O23, 5I75, 6Q0L, 1XIW, 5F8R, 4UV7, 4HZL, 4KJW, 5M94, 6SUZ, 6BKC, 6B0S, 5F3B, 3VRL, 6MLK, 3H3B, 4YBQ, 5U3N, 6X9R, 5K9O, 6D2P, 3J3O, 7A0V, 1XGR, 7D4B, 6DDM, 6BIT, 6U0L, 5LQB, 6BKD, 3WFC, 6DC9, 4KKL, 6X9V, 6WIR, 4KJP, 6B20, 6PV8, 7PIV, 4WUU, 5XJ4, 6ORP, 6H71, 4UTB, 2DQJ, 3V4P, 6RAH, 6EA5, 6MB3, 3ZTJ, 5U6A, 6SNI, 6PLH, 4ZFO, 3ETB, 5KEN, 7K3B, 7LY9, 6O41, 5U4M, 5HGG, 7AEJ, 1U8M, 5NPI, 6OGX, 5NJ3, 4PD4, 7EPB, 6GK8, 7KQG, 7LUE, 6LN2, 5K9Q, 6EY0, 7C2E, 1ZMY, 6Q0E, 6S3D, 3WKM, 7S8L, 1IKF, 5OMN, 4YDI, 4LF3, 5EPM, 7DTY, 6VMS, 6QEE, 6DF1, 7D3L, 2R29, 6XOB, 5T3X, 5VL7, 3WLW, 6MDT, 2DWD, 6X8U, 4WV1, 5UK1, 5KWL, 3WIH, 6VTT, 3EFD, 6OS1, 7O0S, 3O2D, 5FB8, 4YUE, 7L1V, 4LQF, 6UUS, 3P11, 3DVN, 7K5Y, 6MFP, 5E1H, 5GUX, 7F55, 7MBX, 7DR4, 5U3J, 4K94, 5T6L, 4Y5Y, 6WEQ,

6XQ0, 1G7H, 4LMQ, 4HS6, 1C08, 3J8Z, 7NQA, 4JAN, 5TZU, 5B8C, 4HG4, 3ULU, 4MA8, 2P7T, 7DM1, 1YYL, 7DFC, 2JEL, 7B09, 6C08, 6DC3, 4XNU, 4KUC, 5KQV, 6H7J, 1KB9, 4I2X, 2FEC, 4M62, 7KC9, 5D96, 5MZV, 6BQB, 7D3R, 2YC1, 4DN4, 6SNC, 4I18, 6P8M, 6HJY, 5U5F, 7KD6, 5GIS, 3IDY, 3ZLQ, 6RAN, 1UAC, 2I5Y, 4HS8, 5W23, 5I74, 6QTL, 5VCN, 2IH1, 2H32, 5TFW, 4IOF, 4YXL, 5T3Z, 2WZP, 2BRR, 2HTL, 5C0R, 7L0L, 7AQZ, 2CMR, 4KK6, 5WNB, 4NHH, 7KDM, 3TYG, 2XV6, 5VEB, 6Q0O, 6X3V, 4ZFF, 7K63, 6WQO, 5W5Z, 4YX2, 4LHQ, 6VY5, 4LSU, 4LSR, 1V7N, 6VGR, 4I13, 3ZDY, 6OS0, 3MXW, 4KRP, 5HD8, 3WFD, 4DKE, 6LXK, 6O3G, 7DFA, 6CRQ, 6AQ7, 3SN6, 3H42, 6I8S, 1U8N, 3J8D, 6AWQ, 4LDO, 6O1F, 6ETI, 4OD2, 6MTJ, 6FYT, 5I9Q, 6BCK, 6SNE, 5NBD, 6M3B, 6NIU, 6HHD, 6O25, 6P8N, 2R56, 6M0Z, 7LEX, 7EW1, 5OWP, 4XP5, 1G7M, 6T3F, 4Z9K, 5GIR, 2XWT, 6BY2, 5OCK, 6XPZ, 3CSY, 7LBE, 6TYS, 4KXZ, 5GZN, 5W3O, 5VK6, 7MMN, 6U9S, 7F9Y, 7KJI, 7MXE, 6HHC, 6W0G, 4MHH, 2X6M, 6GLW, 4F15, 1FBI, 6AZ2, 6MEK, 6BPC, 6DFH, 4LVN, 6SND, 5VXL, 5DMG, 5JQH, 2R0L, 4KRM, 6VY6, 5NUZ, 3V4V, 6I07, 3H0T, 6RPJ, 3EOA, 7LBF, 2FD6, 6BAN, 6CNV, 3GBN, 1K4C, 6XOX, 3RVV, 3HB3, 4XVT, 6WX2, 6NFV, 6B9Y, 3JWD, 5VCO, 7K7I, 5LHP, 6YYE, 5JW4, 3FFD, 2R0K, 1U91, 7LJA, 6BKB, 5TIH, 6H2Y, 5W0K, 6VN7, 6BFQ, 6PYC, 5HM1, 3TT1, 3N85, 6NCP, 6K69, 4OLY, 3VI4, 7A4T, 6FY2, 1I9R, 5MU0, 6RCU, 3W11, 6VLW, 5O0W, 7NOW, 5CSZ, 6XPR, 4XPA, 5I8C, 4K3J, 1S78, 4JQI, 4P3C, 7NEQ, 7KLC, 4JDT, 2W65, 3U0T, 6DBG, 5D1Q, 6B5N, 5VIY, 4TUK, 2Q8B, 6WHC, 6MQE, 6B0N, 6RPS, 2EIZ, 5T85, 6AVR, 6P50, 5TZ2, 6CSF, 6XP6, 6Y9B, 5F7N, 6F2G, 7K76, 6MEJ, 1UWX, 2FJG, 3JCB, 6B70, 3IFP, 4KKB, 5T29, 3TJE, 3EJZ, 5MVZ, 7KZX, 4FG6, 7CMU, 6K6A, 6X07, 6Z7Z, 4YE4, 5LBS, 6PZW, 7BUF, 5TL5, 6OCB, 5E7F, 5IKC, 6WG0, 6KS0, 2JK5, 3JBQ, 6A4K, 6TNP, 5DLM, 4IDJ, 3A67, 6JFH, 6URM, 4TSB, 6VEP, 6E56, 7JTF, 2P42, 6CEZ, 4NNP, 3VGA, 7S8O, 5TPW, 6QD8, 5D71, 7DWT, 5X2O, 6AWP, 6XZU, 4XZU, 5DS8, 6H15, 4RGM, 5MUB, 4JM2, 7NX3, 3ZDZ, 7BSD, 5SX5, 5O2U, 5WTH, 4RWY, 6E62, 4MHJ, 6DDV, 6MQR, 6Z1V, 4KRO, 6UL4, 7EW7, 7JR7, 6QX3, 6RU5, 7A6E, 6VN0, 7CJ2, 5HI4, 7MLV, 4YQX, 3LH2, 7KEJ, 6IW0, 1TPX, 7EW2, 5YY4, 7A5V, 3MAC, 7RCO, 5DFZ, 4ONF, 6OBM, 3MLV, 7A6F, 6H02, 4FQI, 5KTE, 5DSC, 6TKO, 4MXV, 5YE3, 2QQL, 7NIU, 5FCU, 6MI2, 6II4, 3K1K, 3KLH, 7MFR, 6UMX, 1J5O, 6E3H, 4QXT, 6Z7Y, 5VPL, 1BVK, 4ZFG, 6M58, 4FQJ, 7CFN, 6MID, 2YSS, 7KYL, 6N4B, 1G7J, 6WDT, 5VAQ, 3TT3, 6Z10, 7KF0, 5Y11, 7D5U, 5VKD, 6LPB, 5EC2, 5DHZ, 5NH3, 6NF5, 6VBP, 6EA7, 5KU2, 6RAJ, 4YFL, 7AUE, 6WTU, 1SY6, 4XTR, 1OTU, 3SDY, 6EHG, 6AWO, 3ZKN, 5OTJ, 5I73, 7LKH, 6AL0, 4S1R, 7KGV, 7DOH, 5VGJ, 5HI5, 2J88, 5DHX, 6UTE, 3BN9, 6X05, 5USH, 3V52, 6IR2, 4J6R, 2HT4, 2VXQ, 6O39, 5TUD, 1R3J, 5A7X, 5BVP, 5T5F, 7F8W, 5K59, 1DQJ, 6BDZ, 4DTG, 6LR7, 7P6K,

6UVO, 6AYZ, 6OYZ, 4KK8, 6WZG, 5TE6, 5WKO, 7LFD, 5OCY, 4G7V, 1OB1, 3IDX, 7AR0, 5KVG, 2ZNX, 4HFU, 6OPP, 7F4I, 4KJQ, 6B0E, 7O30, 4G6J, 3P0G, 5M95, 7MFG, 5J1S, 7NCS, 6OEL, 7EC5, 6DCA, 3ULV, 1KIR, 6OMM, 3J70, 7RP0, 4X7C, 3R08, 4QKX, 4YO0, 7AHV, 6HX4, 6L62, 4LSQ, 5USF, 2ZUQ, 6DZV, 4IOI, 1U8I, 7JHH, 1U8P, 6MHG, 3J3P, 2NXY, 5F7K, 6MQS, 4F9P, 6K41, 6YIO, 1VFB, 6IEC, 5X8L, 6JHR, 5GRU, 5VPG, 4DK6, 3D85, 7JVP, 4ZPV, 7DST, 4QYO, 2ADF, 6Q18, 6VYH, 6OY4, 7M8L, 4XMP, 7RK1, 6B3M, 7S6C, 7S13, 6EWB, 6CMO, 3B2V, 7LKF, 6ME1, 6PHF, 6TGG, 6VI4, 4O4Y, 5CJQ, 6QEX, 6NMV, 6FXN, 5CBE, 7E14, 2P48, 3T2N, 6APD, 6PHB, 5MO3, 5C0N, 6H3T, 6IEQ, 5O7P, 7ECY, 7AQY, 4CAU, 6OTC, 1ZTX, 5B71, 1DEE, 6ULE, 3MNW, 5FHC, 2R9H, 7KDD, 6UM7, 7D8B, 3AB0, 6VJT, 3RI5, 1IC5, 5VJ6, 2XQB, 4ERS, 5GS0, 7KH0, 5KW9, 1N5Y, 6AVU, 5JA9, 6NMT, 6EDU, 5X2Q, 6KO5, 6H1F, 6D0X, 3EZJ, 6IDI, 2P45, 7RSN, 3S36, 1NMB, 6PHD, 6GJU, 6U36, 6YXF, 5H8O, 1IQD, 1HYS, 1ZA3, 5UDD, 7M2J, 5X2M, 4TVS, 4XT1, 5TKJ, 6H7N, 4QXU, 6FV0, 3KJ6, 7F16, 4RFN, 6FZQ, 7PHQ, 4F3F, 5BK1, 7NKS, 5YAX, 5GZO, 2P44, 6TYB, 5VIC, 4XXD, 6OCA, 6OQ5, 6P9X, 7D7M, 1U92, 6X4T, 1XGU, 3I50, 6RNK, 4LEO, 5AUM, 6S8D, 6UKJ, 6U52, 7LUC, 7CW0, 6DB6, 4I77, 4HIX, 4X7F, 6CW2, 3MLT, 4OLX, 7KC1, 5D72, 7CFM, 2H9G, 7LF1, 5T80, 6LHQ, 7JKT, 6Q0H, 7LU9, 4I3S, 4BH8, 7DH5, 6PWU, 6NN3, 1YYM, 6VBQ, 3U30, 5VQM, 6BF4, 5U8R, 2VIS, 6MNR, 3HPL, 5VIG, 2DQF, 6RLO, 6CH9, 5VAG, 7DW9, 3HI6, 4JHW, 6WOZ, 6NMS, 3FMG, 4WFH, 5MHS, 6PB0, 7CKZ, 5JSA, 6Z3Q, 5A2L, 7F58, 6LZ9, 4GXU, 3CFI, 6I2G, 6B5R, 3KS0, 6LYN, 5A2J, 6JSZ, 4R4F, 1R3L, 5EII, 5Y9J, 5SV3, 6TOU, 3OGC, 5TOK, 5CIL, 6AWN, 7KDE, 6LFM, 4IRZ, 4FQK, 6UIG, 6UYF, 6II9, 6X3X, 4PP2, 4RRP, 6WNO, 4YGA, 5T1D, 7CKY, 5OCA, 6P9Y, 6MN7, 7BZ2, 4XAW, 5XS7, 7FEI, 1NCB, 6UUD, 6X1V, 6N5E, 6SGE, 6GG0, 1XCT, 2DQH, 6YO6, 1G9M, 7F54, 6QXE, 4KDT, 5W3P, 4PLJ, 6GS1, 1T03, 1GC1, 6KRZ, 6V4N, 6N16, 6KN9, 6DB5, 7C2T, 6VI0, 3J5M, 2NYY, 4DGY, 2B2X, 5CZV, 6PBV, 6E4X, 5U7O, 5KAQ, 1XGT, 4TSC, 1KXQ, 6LNT, 6FN4, 5AAM, 3LIZ, 5XF1, 6WI9, 5XCV, 6MG7, 4G7Y, 3K80, 6QFC, 5YHL, 6HUO, 7S8P, 6P91, 5FHX, 5TD8, 6VX4, 6W2C, 6CWD, 7E9H, 2NZ9, 6ZBV, 4XWG, 4KFZ, 5VTA, 7CU5, 5WTT, 6OOR, 4UT6, 1ORQ, 4GAG, 7A4Y, 5H35, 4V1D, 6QD7, 5U1F, 1CZ8, 3HAE, 6CSE, 5OJM, 7E6U, 3D9A, 4HJJ, 6PWC, 6BFS, 6C5V, 7KFB, 1FE8, 2VC2, 6DFI, 6UUN, 2EXY, 6PDR, 6A78, 3U9P, 5M15, 7C81, 4KKC, 4XNQ, 6XSW, 4OLV, 5EUL, 1FPT, 6RAG, 6ZE1, 7O9S, 6UTA, 4M5Z, 4BFB, 3OGO, 3FB6, 1G9N, 7F2O, 6UTK, 6H6Y, 6X08, 6T3J, 6O3A, 4XCF, 6IBL, 5WB9, 6LI3, 6ORO, 6O3U, 6Q0I, 5KU0, 6WZL, 7BUA, 6CK9, 3L5X, 7KFE, 6XBL, 3LQA, 6VRK, 2HT3, 6GWQ, 6W52, 3UZE, 6X1U, 7CW2, 5WI9, 5XMH, 5U5M, 2DQD, 7MHZ, 5X8M, 5UG0, 5W2B, 4IJ3, 4JPK, 6UM5, 4XC3, 6OLP, 2NY1, 2EXW, 5USL, 6Y90,

3CVH, 6MUI, 7KDU, 4JZJ, 4LGR, 6YXD, 5W1K, 4O5I, 4MQX, 4KKA, 1NCC, 2FEE, 4PS4, 5LHQ, 7LOK, 5WUX, 5H30, 6FRJ, 1MEL, 4MXW, 7CVZ, 5Y2M, 6WO5, 4H88, 5YD4, 6R8X, 4K24, 3SKJ, 3G9A, 4MWF, 7KZZ, 6W16, 5NMV, 7MLU, 7KI6, 5V6L, 7M7G, 7RMH, 5VAN, 6AOD, 7LMR, 6I6J, 5KVF, 5GGV, 7R89, 7CKW, 4LSV, 6NYQ, 6SVL, 3K3Q, 5J13, 4TNV, 5VZY, 6ELU, 6QB3, 1JHL, 3RVW, 4JPW, 7RAI, 5UJZ, 7LG6, 2HVJ, 5I8H, 5DTF, 6OPA, 7M7J, 4TUL, 7KF1, 5FV1, 5VL3, 5MP2, 5F9A, 5KEM, 6K6B, 6MCO, 4NBX, 7O7F, 6KTR, 3RIF, 4K9E, 6CBP, 4QO1, 7MWW, 4OKV, 6OSH, 6WIK, 6OBO, 1A14, 6UDK, 2H2P, 6C9U, 3FB8, 7A48, 7R8B, 6CMI, 5NPH, 6OHG, 6X1S, 7MYZ, 5U3M, 4FFY, 6F7T, 3JBA, 6O42, 4NBZ, 6KZ0, 6DE7, 5VOD, 6P67, 6MTP, 5UQY, 6PEF, 6YWC, 6OZ3, 4JB9, 2HLF, 6LMK, 5GHW, 4ONG, 6S8J, 1YQV, 7KD0, 1JRH, 6CYF, 7K93, 6GWP, 7JWG, 6DBF, 4RQS, 5XEZ, 6NM6, 2DWE, 5LX9, 2ITD, 5F7W, 6UBI, 6BSP, 5UZ7, 6NFC, 1KYO, 7MXL, 6WH9, 6NJL, 5JQ6, 7LSF, 6GKD, 6QX4, 6CF2, 6OZ9, 4HPO, 7F4F, 6XLQ, 6QPG, 3ZE1, 6XZW, 2P47, 6PPG, 7DUO, 6X1W, 1RI8, 6Q20, 6WWC, 4X7E, 2HTK, 4N0Y, 6GJS, 7F4D, 6VRY, 7DNL, 4TVP, 5DD0, 7KEW, 7CZD, 6W09, 6NQD, 6U59, 5GJS, 6FZR, 3T0X, 6JHT, 4XVU, 3W12, 5F7L, 5CJX, 6AXL, 4HJG, 2R4S, 5Y9C, 7CQD, 5UGY, 3ZDX, 5KTZ, 4XVJ, 6YWD, 5AAW, 4EDW, 7EKK, 7BU7, 6GK4, 6XGC, 7O2Z, 7O52, 5JZ7, 4ORZ, 4POU, 6SV2, 2ZCL, 3S88, 2YPV, 4XX1, 4RDQ, 4XNX, 4Q6I, 1XGP, 2NY6, 7DF9, 4JZO, 5N7B, 7EVZ, 5C1M, 5LHR, 7S6D, 5NBM, 3P9W, 4Z7Q, 5ZV3, 5U3K, 4KK5, 6Y92, 2Q8A, 2VDO, 5OB5, 5NJD, 5TLK, 5MO9, 7CKX, 4XI5, 6BPA, 1YNT, 5F9D, 7CX2, 7N0U, 2HVK, 5OVW, 6CDO, 6GFF, 6H7O, 3UC0, 7K37, 4RX4, 5VXR, 3DET, 3FKU, 7LL1, 6D9W, 5O02, 5CUS, 5HBT, 5IBL, 6RAK, 5B3J, 1AHW, 7CZ0, 4TUJ, 6V8Z, 5VKE, 5TE7, 5EOR, 6RCV, 4EJ1, 7R88, 6B5M, 6MTO, 4PGJ, 5MHR, 6MEH, 6GLX, 6X87, 7ND0, 6NBH, 3QA3, 7D85, 6YX9, 6I9I, 3LEV, 5E8E, 1QGC, 5IMM, 1J1X, 5Y80, 3P0Y, 6ZG3, 6XSK, 7MTA, 6HF1, 7JTG, 5ZS0, 5V8L, 5A2K, 5C6T, 4BZ1, 1PKQ, 3RAJ, 6OUS, 4HJ0, 6BF9, 4N1H, 5ESZ, 5T33, 5W3L, 5DUR, 5E1A, 4MSW, 6VJN, 7CBP, 6SC6, 6OKP, 1KIP, 4AG4, 7KR5, 6MUF, 6MJZ, 3IYW, 5D8J, 6RAL, 5I6Z, 1P2C, 5NJ6, 7F53, 5A8H, 6F2W, 3K81, 5VOC, 4DVR, 5ZIA, 1QKZ, 7LBG, 6ZRV, 7M52, 5MEV, 4P2C, 6XKR, 6OYH, 5DFV, 7JKS, 6UM6, 6NZ7, 5TRU, 4N90, 3J6U, 7BUB, 6M2R, 7CVT, 3ZKX, 6M3C, 7EKB, 3JBF, 6QB4, 6U51, 3OR6, 5YE4, 5JDS, 6AXK, 7DUQ, 6FLB, 4C58, 6P9H, 6OFI, 3SQO, 2FJH, 5Y9F, 5YD3, 5HYS, 6X9U, 1XCQ, 4FFZ, 7AQX, 6EJG, 6MZJ, 4LKX, 6JHQ, 5MYK, 2YBR, 6NFJ, 7JHJ, 7BU8, 6V4O, 5UEK, 6U54, 6WF1, 5JA8, 1R3I, 4ZSO, 4XRC, 6MAM, 1R0A, 4G80, 2W0F, 3ZE0, 4JKP, 5D93, 6WJ1, 6OE4, 6BT3, 4GFT, 5MYO, 6IW2, 7DNH, 4NBY, 7C80, 4U1G, 6AZZ, 6BP2, 5N09, 3V7A, 4OLU, 4J4P, 4DKF, 3B2U, 3GI8, 6ZVF, 5IHL, 6OS2, 2H3N, 3BDY, 2OZ4, 4XP1, 6QV2, 5W08, 4WFG, 2QQN, 6P4A, 6CDM, 3EBA, 4DW2, 6P65, 1U8H,

4KHX, 7KYO, 1NBZ, 5MY6, 6K0Y, 3IXY, 3QSK, 6CXY, 5FYL, 7K60, 5V7J, 3F5W, 2EZ0, 1WEJ, 4NRX, 7LWD, 6QGX, 7KET, 6KS1, 4NC1, 6WPW, 6PA0, 5W3M, 4O51, 7LF2, 4I0C, 6B9J, 6VZI, 2LTQ, 3THM, 4YK4, 6UJ9, 6BPE, 5J57, 6PDX, 6K42, 6VN1, 6FE4, 2FED, 6TXZ, 6J5D, 6W0C, 5MYX, 6P3R, 5UHY, 5IQ7, 6Y6C, 7K66, 6OSV, 5T78, 5VPH, 7E7E, 1J1O, 2ARJ, 6K5I, 6ICC, 2VDL, 7DUR, 6SF6, 7KTX, 3CXH, 5MI0, 1E6J, 6LBH, 6BF7, 6CE0, 6WFZ, 5E94, 4M1G, 6BLH, 5DYO, 7KF9, 2Y6S, 6MV5, 4L5F, 5MP3, 5T5N, 4HWB, 2VH5, 4XH2, 6CH8, 2IFF, 6B5O, 4CDG, 2IBZ, 4QWW, 6GS7, 1FDL, 1R3K, 4Z0X, 4WFE, 3FB5, 5GXB, 6CNJ, 6A96, 6ID4, 1JTP, 3UZQ, 7DHR, 7E9G, 6JBT, 5C0S, 6AZM, 6N7J, 5IP4, 6S3T, 6DDR, 7EBR, 1OTT, 6HUG, 7ANQ, 6XY2, 6LGW, 6H5N, 1A3R, 7BU6, 6UVA, 6UYE, 4CMH, 6R7T, 7MN8, 6U0N, 7JHG, 6Q6Z, 5C2U, 7C2S, 4U3X, 6CHB, 3IXX, 2VDM, 4D9Q, 6U2F, 5LCV, 6FLC, 6A67, 4Y5V, 7LSG, 1A2Y, 1QLE, 5IWL, 6WAS, 7JWP, 5VN3, 6GK7, 6LRA, 6ADC, 4J8R, 4LST, 6MQM, 7BBG, 3IU3, 1ZV5, 5D70, 6XW4, 3ZTN, 5HVF, 4RIS, 7JJO, 5ANY, 6AD0, 5U3L, 6P62, 6OIK, 7AQG, 7A50, 6PHC, 7A65, 3MJ9, 6WBV, 6NHA, 3V4U, 3RU8, 6YSQ, 6CBV, 6BZW, 6ITC, 6NBI, 1NCD, 5C7X, 3ZKQ, 6LHT, 6N81, 6CRK, 1QNZ, 6JOD, 5CJS, 4OII, 6WZJ, 7BUE, 7BB7, 5FOJ, 5MP5, 5SY8, 4MQT, 4K3H, 5ERW, 4P9H, 6IDK, 6HUJ, 6M1H, 6X3T, 4MA7, 3RIA, 4Y5X, 5TZT, 5M30, 6OAN, 6CO3, 6X1A, 5VAK, 7CE2, 3IXT, 4PIR, 6XQW, 4XP6, 7BQ5, 5MV3, 2NY0, 1NDM, 1CT8, 6PBW, 6ION, 7AHU, 3O41, 6ATT, 7F4H, 2HJF, 4JZN, 4OM1, 5XXY, 6WFX, 6W0E, 2NY2, 6H16, 7RAH, 7LMQ, 4F9L, 6SC7, 6MNQ, 7D5P, 5F9W, 5W6G, 7NX0, 7KFH, 6V7Z, 6UQR, 1ZVH, 1G7L, 4BH7, 6K3M, 6WFY, 7LFA, 7M7I, 3K7U, 1JTT, 5BOP, 7C61, 7NW3, 3IGA, 3J8V, 7MHY, 2P43, 5HJ3, 4HEP, 6VY2, 6VEL, 5D1X, 6M1I, 5UDC, 7DFL, 6PZE, 7A17, 3JCX, 4QTI, 5XCR, 2NR6, 4CNI, 6C9W, 3T0W, 4XNM, 5T6P, 6TYL, 4RFO, 4I1N, 6BA5, 6P7H, 3QUM, 5OGI, 6K64, 5LDN, 6I8H, 6PEC, 7KQH, 6OBG, 2VDQ, 4YR6, 7N0V, 1W72, 7V9M, 4LDE, 6HXW, 4G6F, 5N88, 3NIF, 3GJF, 6O8D, 3NFP, 5GS2, 6OZ4, 6USF, 7LDE, 4FQV, 2BOC, 6P4B, 6KVA, 6MTQ, 5DHY, 1EZV, 3RVX, 5IQ9, 7EO0, 4W2O, 6MWV, 6O26, 4LHJ, 5W1M, 6B0G, 3MLW, 6PLK, 3ZKS, 3W2D, 6XZF, 4YWG, 6WG2, 6W05, 6WOQ, 6H7L, 4Y7M, 7DWU, 6X3S, 6BFT, 1KEN, 5EN2, 5EBL, 4OCN, 7NIW, 3STZ, 5ZUD, 6D6T, 6PT0, 6GJQ, 6NJM, 6BZU, 5IML, 6OPO, 4OT1, 5O05, 6Q23, 5I5K, 1XF5, 3S37, 3MA9, 7DFP, 5UCB, 6WW5, 5UMI, 2VIR, 5NJG, 5VNW, 5MP1, 6QNO, 6W2B, 6JFI, 6QGW, 1P84, 4UAO, 5UK2, 7BW0, 4KI5, 1NCA, 6JB8, 4M48, 4BZ2, 3JWO, 5L6Y, 5EZO, 1FSK, 2R6P, 6B5S, 4YDL, 6OQ6, 6WGL, 6B0H, 6DC8, 6N48, 5W42, 5UMN, 4FQY, 6E0P, 6MTT, 7DSS, 1OTS, 4YJZ, 6E4Y, 2R69, 1UA6, 3PP4, 4Z7N, 6QB6, 5LXG, 4XP9, 7FEJ, 5O14, 3CX5, 3NCY, 6FLA, 7MEM, 4YDJ, 1TQB, 5TJW, 7LVW, 5J9P, 5G5R, 6VCB, 2B4C, 7KMD, 1ORS, 5MES, 5X08, 7EVY, 5Y7Z, 2R4R, 7BSC, 6HJP, 6N8D, 2DQI, 4DKA,

6ORN, 3LDB, 3UZV, 4NZT, 3T3P, 3C09, 4PLK, 5W5X, 5DHV, 4G3Y, 3VI3, 7EAK, 2VIT, 6N5B, 6H0E, 4WFF, 5KN5, 6SC5, 3JAB, 5EC1, 6E5P, 7KI4, 6NFU, 6QUP, 1FJ1, 7LMN, 1YY9, 6LDY, 6LDV, 6RVC, 5O1R, 4XVS, 1JNH, 6SSI, 6AJ9, 5LWY, 5F72, 6O29, 1K4D, 6QGY, 7NJZ, 5O04, 5F6J, 2HT2, 2X89, 6YE3, 5Y0A, 6IUV, 1IC7, 5V8M, 3EOB, 6B5T, 5UK4, 3R1G, 7SEG, 6PCI, 7NIV, 7EW0, 5IMK, 7M3N, 4S1Q, 6CVK, 1UJ3, 6RTW, 6J6Y, 6X9T, 6UR5, 7M2H, 1MLC, 6D11, 2JIX, 4H8W, 4TQE, 5GGT, 7MTB, 5BK2, 6K68, 6BZV, 6OQ7, 6VO0, 3BSZ, 6SC9, 5HVH, 6CMG, 6VYV, 5TE4, 3HFM, 6PI7, 3U2S, 6LZ2, 6XW7, 7O33, 4WGW, 6VBG, 2QQK, 6OYA, 6P95, 6H7M, 5L0Q, 4YZF, 5SX4, 6OBC, 4WHY, 7LJB, 2Y5T, 4S1D, 5XHV, 4LVH, 7LEY, 3MLX, 5FUC, 7CX3, 5I71, 1BQL, 7S8N, 6U3I, 6M47, 5WN9, 6UFT, 5E2V, 3UAJ, 2XRA, 7E72, 6O3L, 1KXT, 1OAK, 2Y07, 3JBD, 4TNW, 6VLN, 4U6V, 5IFJ, 7JTR, 6AD7, 3G6D, 6U1N, 6H06, 6PCU, 4UTA, 7K7H, 6WM9, 4C59, 7DFB, 5TR1, 5DRZ, 4W6Y, 6NJN, 6X3U, 6W0I, 6W0F, 6UKT, 5VOB, 3UO1, 6CDP, 7A6C, 5FEC, 1QFU, 3NIG, 6DFG, 3WFE, 4EIG, 3EJY, 6XBJ, 5WDF, 4XCY, 1XFP, 7KBB, 7R8A, 5BK0, 4NC2, 1NDG, 7BB6, 5JS9, 4HPY, 5DUB, 6DB7, 5O4G, 6PV7, 6Z2L, 6CM3, 3VE0, 6DDE, 5VXJ, 7DQ7, 3IET, 4OM0, 5UTF, 6HHU, 4P3D, 4WEU, 6WVZ, 5HDQ, 6A79, 1U8J, 6WHK, 7EB2, 7K7Y, 6WMW, 7CW3, 6IR1, 6D01, 5EA0, 4R4N, 5YWP, 5ZUF, 3JAU, 5OCX, 6FEQ, 6W0B, 6VCA, 4NZR, 3K74, 3L95, 6NIP, 4ETQ, 7CX4, 5UTZ, 5A3I, 3MOD, 6WZT, 6IUT, 6EK2, 6NMR, 6GS4, 6W03, 2XT1, 5G5X, 6PDS, 4BKL, 7CVY, 6LDW, 6Z06, 7M51, 5EU7, 6WIX, 6MUG, 6B5L, 6ADB, 6U6U, 4X7D, 3V0A, 6SNH, 6IBB, 7P5W, 5HDB, 5NQW, 6ARU, 5NGV, 1U8K, 2ATK, 7LJ4, 5T5B, 4R3S, 4OCL, 7DNK, 7KQI, 3O0R, 1RZJ, 6EQI, 6HJX, 6OPQ, 2VDK, 4QNP, 7BTS, 6ULC, 6X8Q, 4C57, 4YXK, 3ZKM, 7DGE, 7EPU, 6WF0, 4NM8, 6MVL, 2J6E, 3NPS, 7JV5, 6FY3, 1BZQ, 6O28, 2UUD, 6MNS, 6U55, 6ZPL, 3KR3, 6WO3, 6K5D, 7S8M, 4KML, 5UEA, 4ZPT, 7D5B, 4YHP, 6GV4, 6J15, 7LDD, 6HUP, 4W6W, 4OQT, 5H37, 4GAJ, 4LVO, 6XXV, 4KK9, 7K61, 5W0D, 6O2C, 6OCD, 6FUZ, 2NY5, 6XSN, 6PZF, 6Q1Z, 6UCE, 7JIE, 7KBK, 6OE5, 5E0Q, 4Z7O, 6X3W, 4QEX, 6XPQ, 4WY7, 3ZE2, 2EKS, 1MVF, 7LF8, 5TLJ, 7OH1, 1BJ1, 4WEN, 6X19, 4AM0, 3NID, 6NBF, 6W51, 6FYW, 5J3D, 5OCC, 4AL8, 4LCU, 6UI1, 6W0H, 6LHP, 6MU8, 1RZK, 2I60, 6CH7, 4JPV, 7LJ5, 7CVS, 7KCR, 6K5A, 5TQ2, 6OAO, 4OLZ, 6RAF, 6WER, 7DAA, 5L21, 2XTJ, 7VUX, 5VK2, 6E63, 6MU6, 7DHI, 4LU5, 6APP, 6V6W, 7P60, 4UIF, 3STL, 6XLI, 5GRJ, 6XW6, 6GWN, 6HCO, 7JVR, 7F9Z, 5WT9, 6B73, 4R8W, 1OAZ, 1J1P, 3QWO, 7CHY, 3SOB, 6BLI, 7LFB, 6O2A, 5F9O, 6B5P, 6BZY, 4RAV, 4JO1, 4K8R, 7C6A, 3K2U, 7EXD, 3PNW, 7LJC, 7RNN, 6CXG, 6UHT, 7JOO, 4DK3, 6AJ7, 4Q0X, 7JWQ, 7KPG, 4P59, 5IGX, 5BOZ, 5UKR, 4TSA, 6VMJ, 1JPS, 5JXE, 6S8I, 6E0C, 6XQ2, 6O9H, 7M7F, 6WIT, 7KJH, 3JBC, 5CJO, 2NLJ, 6I8G, 4OGA, 1LK3, 3SE9, 3STB, 6JB5, 3Q1S, 5KVE, 1NBY,

1TZI, 7PHP, 6EYO, 5THR, 7KD2, 5IJK, 4OGX, 4KVN, 6N4Q, 6KX1, 7D77, 4K2U, 7JTI, 6VRH, 5JMO, 4XAK, 5JYL, 7A4D, 7K5X, 5DUP, 3P30, 7NRH, 7LL2, 4FFW, 4DQO, 4M3K, 6AL1, 6U12, 6XQ4, 5HBV, 7CHZ, 7DB6, 7AZB, 5KOV, 6KPF, 5CEZ, 6C5W, 2ZJS, 3W9E, 7KI1, 1EGJ, 6UMG, 3B9K, 6DCW, 5IMO, 6N6B, 6JB2, 5VJQ, 2J5L, 1G6V, 2QAD, 6OBE, 1AFV, 5J1T, 6J5F, 5F3J, 5TKK, 6I04, 7D68, 6AVQ, 6A3W, 1KB5, 6ML8, 4GRW, 7E53, 5VYF, 6OEJ, 1DZB, 6B08, 5Y2L, 7L5J, 4YPG, 6RAM, 4XPH, 1TQC, 5O4O, 6PDU, 7E32, 5KJR, 7DHA, 3UJJ, 4PP1, 6OKN, 6UCF, 6FY0, 6VKN, 6XPX, 6A77, 4O9H, 4KRL, 4G6M, 4ZYP, 5W3E, 1G7I, 3RKD, 5NPJ, 6OT0, 3CK0, 1ACY, 1NMC, 6CW3, 5VAI, 5UKB, 4JR9, 4YDK, 7MJS, 5E2W, 4EDX, 4WEB, 3JCC, 6DID, 5HHV, 4RGN, 7PIJ, 6OC3, 2ZCH, 6LDX, 7A69, 6BY3, 2UZI, 6IDL, 6X04, 5TOJ, 7EVW, 7RXC, 3RJQ, 5EBW, 6GKU, 5X0T, 6GCI, 6BB4, 6JHS, 5JYM, 2W9E, 5WOB, 5M2J, 2IH3, 5HVG, 6OQ8, 5D1Z, 6FAX, 5XJ3, 5C8J, 6E4Z, 6PIS, 7EW3, 7LJD, 5M2M, 1U8Q, 6WTV, 6VMK, 7CRH, 6DZL, 1XGQ, 4UT9, 6PB1, 6QFA, 7C4S, 6VEQ, 4XNY, 7FIG, 5UTY, 7P5V, 6YXK, 6O3J, 6IEA, 6VY4, 4U0R, 6B7Z, 6P60, 5JW3, 3UYR, 7CN2, 5M2I, 7NGH, 7RK2, 7RMG, 3SE8, 5CWS, 7S11, 4HC1, 7BXV, 3EHB, 4JO2, 4OJF, 2WUC, 6KNM, 5O8F, 7JUM, 4QCI, 4FFV, 1NSN, 4TTD, 6VO1, 2BOB, 3A6C, 5NBL, 6ICF, 5GJT, 5LXA, 6U02, 7LLK, 4XPF, 3WD5, 5UOE, 6QIG, 7JOZ, 6HIG, 2I9L, 4S10, 6AL5, 5X2N, 2KH2, 5KEL, 4NIK, 5C7K, 7BXA, 6OBD, 2AEQ, 6FN1, 1FNS, 6WTY, 6AD8, 5XJM, 6X5B, 3Q3G, 6UYM, 4XMM, 5VKH, 3DVG, 6OGE, 5VJO, 6KYZ, 3LZF, 5MWN, 2XQY, 6H3U, 6W00, 5MV4, 7KPB, 4D3C, 5U8Q, 6UJB, 4YC2, 4R2G, 6X9S, 5EOQ, 7D3M, 4LBE, 6W4S, 6KPG, 7EZH, 6YXG, 6P8D, 4HF5, 7P5Y, 3X3F, 3UX9, 5YWO, 6DZZ, 7KEZ, 7OH0, 1RJC, 4N1E, 1NFD, 3FB7, 6Z6V, 6W1C, 2HFG, 5D9Q, 6W0J, 4UBD, 5V6M, 3G6J, 5TQQ, 6CCB, 4RGO, 5EBM, 2ZCK, 6VBO, 6CT7, 6RTY, 1YMH, 4F2M, 4LOU, 5F7M, 6DZM, 3OPZ, 6DC5, 6YU8, 6UYG, 5KAN, 1A8J, 4OLW, 4CKD, 7K65, 1JTO, 6VVU, 6X8S, 1H0D, 4LSS, 7DM2, 6ZWK, 6RP8, 4YHZ, 6WG1, 6UDA, 2BSE, 7L1U, 6O2B, 6RAI, 5J56, 7LSE, 7KRA, 4IOS, 3RHW, 4ODX, 5TQ0, 6NNF, 6Y9A, 1U8O, 5CBA, 7DC8, 5VXM, 6EQC, 5FGC, 7EBZ, 3F7Y, 6E3Y, 6XW5, 6IVZ, 3J8W, 4JRE, 3V6O, 4FHB, 6OS9, 3WFB, 5GGS, 6MWX, 3W13, 4HKX, 4YBL, 6UYN, 6UL6, 7K3A, 6MFT, 6R0X, 6XCJ, 5FV2, 4LIQ, 5J3H, 1OSP, 1OP9, 4R4H, 6LXJ, 7D5Z, 6B9Z, 7BUD, 6N4R, 5KVD, 6MYY, 4N8C, 6PZY, 2DQE, 4FP8, 6MEI, 4I9W, 6X78, 5DMJ, 5U7M

**Supplementary Table S3.** Results of the EMoMiS pipeline forward phase for motifs with significant structural alignment RMSD score (p-value<0.1). The surface accessibility and DL binding scores were not filtered.

| PDB Spike | PDB database | Sequence match | Database ag-contact | Database ab-contact | Spike ag-contact | RMSD | Struct p-val | DL score | DL p-val | Surface accessible |
|---|---|---|---|---|---|---|---|---|---|---|
| 7L2F | 1V7N | TQLPP | Z:111,Z:112,Z:113 | K:33,K:101,O:48 | A:23,A:24,A:25 | 0.9 | 0.093 | 1.281 | 0.010 | TRUE |
| 7K43 | 6PE8 | ESEF | U:65 | A:53 | E:155 | 0.42 | 0.095 | 1.749 | 0.032 | TRUE |
| 7VNE | 6FN1 | NITN + | A:91 | C:59 | B:332 | 0.41 | 0.092 | 2.259 | 0.092 | FALSE |
| 7M6I | 6WEQ | NLVK | D:345,D:346 | L:34,L:32 | B:533,B:534 | 0.25 | 0.061 | 2.347 | 0.108 | TRUE |
| 7P7A | 6XCJ | E I+R NITN | G:278 | L:99 | E:333 | 0.43 | 0.097 | 2.651 | 0.178 | TRUE |
| 7K8X | 7NWL | N+ L GEVFN | B:210 | D:58 | B:340 | 0.33 | 0.021 | 2.798 | 0.219 | TRUE |
| 6ZXN | 6WEQ | K NLVK | D:345,D:346 | L:34,L:32 | B:533,B:534 | 0.38 | 0.086 | 3.021 | 0.293 | TRUE |
| 7M53 | 7M51 | FKEELD F | A:1234,A:1236,A:1237 | H:104,H:61,L:100 | A:1149,A:1151,A:1152 | 0.09 | 0.004 | 3.057 | 0.306 | TRUE |
| 7M53 | 5UDC | K+ELDKY | F:83 | H:99 | A:1152 | 0.27 | 0.018 | 3.494 | 0.477 | TRUE |
| 7V26 | 5MY6 | HKNN | A:175 | B:101 | C:147 | 0.24 | 0.059 | 3.757 | 0.584 | TRUE |
| 7FG2 | 5NHR | FNCY | D:83 | B:95 | A:487 | 0.3 | 0.070 | 4.034 | 0.691 | FALSE |
| 7L2D | 5CWS | I L+ ILSR D | K:175 | H:54 | A:982 | 0.35 | 0.079 | 4.121 | 0.722 | FALSE |
| 7P79 | 6PDX | GYFKI S I | C:257 | O:53 | A:200 | 0.88 | 0.089 | 4.825 | 0.905 | FALSE |
| 7M53 | 3MNW | ELDK | P:664 | A:30 | A:1153 | 0.14 | 0.045 | 5.600 | 0.982 | TRUE |
| 7M53 | 7AEJ | ELDK | A:664 | D:56 | A:1153 | 0.42 | 0.095 | 8.240 | 1.000 | TRUE |
| 7KQE | 6NC2 | AGST | N:528 | S:100 | A:477 | 0.27 | 0.064 | inf | 1.000 | TRUE |
| 7JW0 | 5W42 | GYFKI S I | A:259 | L:27 | E:202 | 0.92 | 0.097 | inf | 1.000 | FALSE |
| 7L58 | 3B2V | ILPV | A:348 | H:55 | B:727 | 0.38 | 0.086 | inf | 1.000 | FALSE |

**Supplementary Table S4.** Results of the EMoMiS pipeline reverse phase for motifs with significant structural alignment RMSD score (p-value<0.1). The surface accessibility and DL binding scores were not filtered.

| PDB Spike | PDB database | Sequence match | Spike Ag contact | Spike Ab contact | Database Ag | RMSD | Struct p-val | DL score | DL p-val | DB-Ag Surface accessible |
|---|---|---|---|---|---|---|---|---|---|---|
| 7K8X | 5I5K | PGQT | C:414 | F:52 | A:586 | 0.39 | 0.088 | 1.3956 | 0.0135 | FALSE |
| 7L2C | 4Z7N | GDSS | B:253 | C:1 | D:76 | 0.26 | 0.063 | 1.7048 | 0.029 | TRUE |
| 7RW2 | 6BT3 | KHTP | C:207 | I:100 | K:431 | 0.21 | 0.055 | 1.8285 | 0.0384 | TRUE |
| 7KQE | 6GFF | LYNS | C:369 | J:100 | C:81 | 0.29 | 0.068 | 1.8969 | 0.0446 | FALSE |
| 7N0G | 6QEE | GKIA | C:417 | Z:105 | A:894 | 0.38 | 0.086 | 1.9018 | 0.0451 | FALSE |
| 7N9B | 6ULC | PGQT D | B:506 | F:1 | A:315 | 0.43 | 0.097 | 1.9991 | 0.0554 | FALSE |
| 7NDA | 6PWU | NITN | A:333 | H:2 | E:140 | 0.37 | 0.083 | 2.0117 | 0.0568 | TRUE |
| 7N4M | 6GFF | LYNS | A:369 | H:102 | C:81 | 0.25 | 0.061 | 2.0614 | 0.0629 | FALSE |
| 7RNJ | 7M51 | FKEELD FK | B:1149,B:1151,B:1152 | H:101,L:94,H:33 | A:1234,A:1236,A:1237 | 0.1 | 0.004 | 2.1299 | 0.0721 | TRUE |
| 7S3N | 7M51 | FKEELD F | A:1149,A:1151,A:1152 | L:33,H:61,L:101 | A:1234,A:1236,A:1237 | 0.11 | 0.004 | 2.153 | 0.0754 | TRUE |
| 7JMW | 6PV7 | DDFT | A:428,A:429 | H:97,H:97 | E:173,E:174 | 0.31 | 0.071 | 2.1544 | 0.0756 | FALSE |
| 7ND4 | 7A6E | GKIA | C:417 | L:51 | A:895 | 0.3 | 0.07 | 2.4573 | 0.1304 | TRUE |
| 7CR5 | 7S6C | GTTL | A:166 | L:98 | A:350 | 0.28 | 0.066 | 2.5189 | 0.1443 | TRUE |
| 6ZDH | 6OQ5 | DFTG I | C:429,C:430 | I:92,I:92 | A:1926,A:1927 | 0.28 | 0.066 | 2.5368 | 0.1485 | TRUE |
| 7N5H | 6ULC | PGQT | C:414 | K:56 | A:315 | 0.36 | 0.081 | 2.576 | 0.1581 | FALSE |
| 7S3N | 5XEZ | K ELDK | A:1152,A:1153 | L:101,H:109 | A:1046,A:1047 | 0.18 | 0.05 | 2.8985 | 0.2511 | TRUE |
| 7L2D | 6SSP | TPGD | A:252 | H:101 | G:52 | 0.26 | 0.063 | 2.9996 | 0.2854 | FALSE |
| 7K90 | 5OGI | GVEG | C:483,C:484 | O:58,O:56 | A:187,A:188 | 0.34 | 0.077 | 3.1272 | 0.3316 | TRUE |
| 7VND | 7KBT | L SKVG | C:444 | V:45 | A:1992 | 0.34 | 0.077 | 3.3982 | 0.4377 | TRUE |
| 7N0R | 7DNH | NNAA | B:154 | D:100 | A:179 | 0.3 | 0.07 | 3.4349 | 0.4527 | TRUE |
| 6WPS | 6QEE | NITN | E:333 | F:30 | A:92 | 0.41 | 0.092 | 3.6559 | 0.5432 | TRUE |
| 7L2D | 6SSP | TPGD | A:252 | H:101 | B:52 | 0.31 | 0.071 | 3.9322 | 0.6528 | TRUE |
| 7N9E | 2NZ9 | PPEA | B:987 | D:107 | A:63 | 0.39 | 0.088 | 4.2346 | 0.7593 | TRUE |
| 7VNB | 5LDN | GVEG + + +P | B:483 | A:56 | A:187 | 0.41 | 0.092 | 4.3917 | 0.8067 | TRUE |
| 7K8Z | 7OH0 | NNLD YN | C:440 | M:55 | A:1220 | 0.34 | 0.077 | 4.6197 | 0.8643 | TRUE |
| 7N3C | 7S6D | TEGA | C:136,C:137 | H:105,H:103 | A:617,A:618 | 0.12 | 0.042 | 6.5442 | 0.9989 | FALSE |
| 7MJI | 6EQC | GVEG + + +P | B:483,B:484 | E:82,E:79 | C:187,C:188 | 0.43 | 0.097 | inf | 1 | TRUE |
| 7V26 | 6QPG | VGGN | E:446,E:447 | f:95,e:60 | H:274,H:275 | 0.42 | 0.095 | inf | 1 | FALSE |
| 7NTC | 6GFF | F STEK | C:96 | L:51 | D:290 | 0.32 | 0.073 | inf | 1 | TRUE |
| 7L09 | 6LN2 | QDSL | A:937 | M:99 | A:223 | 0.17 | 0.049 | inf | 1 | TRUE |
